# An NlpC/P60 protein catalyzes a key step in peptidoglycan recycling at the intersection of energy recovery, cell division and immune evasion in the intracellular pathogen *Chlamydia trachomatis*

**DOI:** 10.1101/2022.11.17.516864

**Authors:** Jula Reuter, Christian Otten, Nicolas Jacquier, Junghoon Lee, Dominique Mengin-Lecreulx, Iris Löckener, Robert Kluj, Christoph Mayer, Federico Corona, Julia Dannenberg, Sébastien Aeby, Henrike Bühl, Gilbert Greub, Waldemar Vollmer, Scot P. Ouellette, Tanja Schneider, Beate Henrichfreise

**Affiliations:** Institute for Pharmaceutical Microbiology, University Hospital Bonn, University of Bonn, Bonn, Germany; Institute of Microbiology, University Hospital Center and University of Lausanne, Lausanne, Switzerland; Department of Pathology and Microbiology, College of Medicine, University of Nebraska Medical Center, Omaha, Nebraska, USA; Université Paris-Saclay, CEA, CNRS, Institute for Integrative Biology of the Cell (I2BC), Gif-sur-Yvette, France; Interfaculty Institute of Microbiology and Infection Medicine, Organismic Interactions/Glycobiology, Eberhard Karls Universität Tübingen, Tübingen, Germany; Centre for Bacterial Cell Biology, Biosciences Institute, Newcastle University, Newcastle Upon Tyne, UK

## Abstract

The obligate intracellular *Chlamydiaceae* do not need to resist osmotic challenges and thus lost their cell wall in the course of evolution. Nevertheless, these pathogens maintain a rudimentary peptidoglycan machinery for cell division. They build a transient peptidoglycan ring, which is remodeled during the process of cell division and degraded afterwards. Uncontrolled degradation of peptidoglycan poses risks to the chlamydial cell, as essential building blocks might get lost or trigger host immune response upon release into the host cell. Here, we provide evidence that a primordial enzyme class prevents energy intensive *de novo* synthesis and uncontrolled release of immunogenic peptidoglycan subunits in *Chlamydia trachomatis*. Our data indicate that the homolog of a *Bacillus* NlpC/P60 protein is widely conserved among *Chlamydiales*. We show that the enzyme is tailored to hydrolyze peptidoglycan-derived peptides, does not interfere with peptidoglycan precursor biosynthesis, and is targeted by cysteine protease inhibitors *in vitro* and in cell culture. The peptidase plays a key role in the underexplored process of chlamydial peptidoglycan recycling. Our study suggests that chlamydiae orchestrate a closed-loop system of peptidoglycan ring biosynthesis, remodeling, and recycling to support cell division and maintain long-term residence inside the host. Operating at the intersection of energy recovery, cell division and immune evasion, the peptidoglycan recycling NlpC/P60 peptidase could be a promising target for the development of drugs that combine features of classical antibiotics and anti-virulence drugs.

**Author Summary:** Free-living bacteria are wrapped by a peptidoglycan cell wall. This envelope provides shape and protects from osmotic stress, while also triggering human immune defense. Chlamydial pathogens live inside human cells and dispensed with a cell wall in their isotonic niche. Instead, they produce a transient peptidoglycan ring that is essentially involved in cell division. Recycling of peptidoglycan is likely important for chlamydiae to maintain a complete cycle of peptidoglycan ring synthesis and cell division. Moreover, the process may help to save energy and to protect from recognition through the immune system. Despite a potentially central role in cell viability and pathogenicity almost nothing is known about recycling of peptidoglycan in chlamydiae. The minimalist organisms lack all enzymes known to catalyze the process in model organism *Escherichia coli*. Here, we found an NlpC/P60 enzyme to serve a critical step in decomposing peptidoglycan ring derived peptides in *Chlamydia trachomatis*. Our data indicate that the peptidase recycles energy cost intensive components and breaks down the minimal recognition motif of innate immune factor NOD1. We also identified a natural lead compound to inhibit the NlpC/P60 enzyme opening the way for innovative anti-chlamydial drug development at the intersection of energy recovery, cell division, and immune evasion.

## Introduction

*Chlamydiaceae* are human pathogens of major public health concern. These Gram-negative bacteria cause ocular, respiratory, and sexually transmitted diseases and replicate exclusively inside eukaryotic host cells. In the course of coevolution with their host, chlamydial genomes were streamlined to a minimum set of genes needed for their obligate intracellular lifestyle [1]. Inside their intracellular niche, *Chlamydiaceae* are protected from osmotic challenges and the reductive adaptation to their host is reflected by the loss of an energy cost-intensive peptidoglycan (PGN)-based cell wall that wraps free-living bacteria to provide mechanical strength and osmotic stabilization [2–4]. However, the pathogens produce a transient PGN ring to support cell division [5,6].

PGN is unique to bacteria and consists of a meshwork of linear glycan chains connected by crosslinked peptides. PGN biosynthesis begins in the cytoplasm with the formation of UDP-*N*-acetylglucosamine (UDP-GlcNAc) and UDP-*N*-acetylmuramyl(UDP-MurNAc)-pentapeptide. These precursors are used to build lipid II at the inner leaflet of the cytoplasmic membrane [7]. Lipid II is then translocated to the periplasmic side of the membrane and incorporated into the PGN meshwork [7,8]. Glycosyltransferases polymerize the glycan chains from lipid II into long glycan strands and transpeptidases crosslink the peptide side chains [7].

*Chlamydiaceae* have a unique minimal cell division machinery which lacks the tubulin-like proteinFtsZ, the central organizer of cell division in almost all bacteria, and instead use the actin-like protein MreB that is typically used for cell elongation in rod-shaped bacteria [9–13]. It is not fully understood how MreB orchestrates the tightly interlinked chlamydial machineries for PGN ring biosynthesis and cell division. In addition to MreB and its regulator RodZ [13], *Chlamydiaceae* have conserved a reduced, yet complete pathway for lipid II biosynthesis, including unusual enzymes and routes to synthesize components such as D-amino acids and *meso*-diaminopimelic acid (mDAP), the SEDS proteins FtsW and RodA, which are implicated in glycosyltransferase activity and lipid II translocation, and the lipid II flippase MurJ [14–17]. Like other intracellular bacteria with a reduced PGN structure such as *Wolbachia* and *Orientia, Chlamydiaceae* harbor a small set of penicillin-binding proteins (PBP) comprising two homologs of monofunctional PGN transpeptidases (PBP2 and PBP3) which are likely involved in the incorporation of lipid II, and DD-carboxypeptidase PBP6 [10,18,19]. In addition, the reduced-genome organisms retained one homolog of cell division amidases (AmiA) [20] in addition to the cell envelope-spanning Tol-Pal complex [21]. The unusual process of cell division in *Chlamydiaceae* is initiated by a budding mechanism similar to that in FtsZ lacking Planctomycetes [22]. The asymmetric cell poles then mature into two approximately equally-sized daughter cells separated by an MreB-controlled septum containing a transient PGN ring which is remodeled during constriction and degraded afterwards [5,23].

In free living bacteria, PGN remodeling and degradation during cell growth and division is accompanied by continuous PGN biosynthesis, which requires a steady supply with PGN precursors. In addition to *de novo* PGN synthesis, bacteria developed efficient recycling machineries to re-use the fragments released during PGN turnover. In the model organism *E. coli* up to 40 % of the existing PGN is turned over in every generation and 90 % of this material is recycled and reenters lipid II biosynthesis [24,25]. PGN turnover products can enter the cell either as anhydromuropeptide (GlcNAc-1,6-anhydro-MurNAc-tetrapeptide) transported by major facilitator superfamily transporter AmpG or as free peptides released by *N*-acetylmuramyl-L-alanine amidases and transported by the oligopeptide permease complex OppABCDF [25– 28]. After uptake by AmpG the anhydromuropeptide is further processed by NagZ and AmpD yielding GlcNAc, 1,6-anhydro-MurNAc and the free tetrapeptide L-Ala-γ-D-Glu-mDAP-D-Ala. LD-carboxypeptidase LdcA removes the terminal D-Ala which together with the tripeptide reenters the biosynthesis pathway [25,27]. The tripeptide is either directly reutilized by UDP-MurNAc:tripeptide ligase Mpl or further processed by MpaA and YcjG. MpaA hydrolyzes the non-canonical γ-D-Glu-mDAP peptide bond within the tripeptide releasing mDAP from L-Ala-D-Glu. YcjG catalyzes epimerization of L-Ala-D-Glu to L-Ala-L-Glu [25,27].

In the highly streamlined *Chlamydiaceae*, recycling of PGN turnover components may not only be important to minimize loss of energy cost-intensive intermediates, but also to maintain a full cycle of PGN-ring biosynthesis, including remodeling and degradation, which is essentially interlinked with the process of cell division. Moreover, PGN turnover products are so-called pathogen-associated molecular patterns (PAMPs) and *Chlamydiaceae* need to tightly control the pool of immunostimulatory PAMPs to evade eradication by the immune system and to sustain long-term residence inside their host. Thus, recycling of PGN fragments may also be a crucial pathogenic strategy to subvert host immunity. Despite the potential importance of PGN recycling for chlamydial biology, almost nothing is known about the process and its impact on chlamydial biology. So far, several chlamydial PGN remodeling and degrading enzymes were identified to contribute to the release of PGN fragments, i.e. dual functioning lytic transglycosylase and muramidase SpoIID, which hydrolyze the glycan backbone of PGN, and DD-carboxypeptidases PBP6 and LysM protein Cpn0902, which remove terminal D-Ala from the pentapeptide side chain [18,20,29]. In addition, we found *Chlamydia pneumoniae* AmiA to act both as DD-carboxypeptidase and amidase, releasing the peptide side chain from the sugar moiety [20]. Recently, *C. trachomatis* was shown to encode the oligopeptide permease complex OppABCDF facilitating transport of PGN-derived peptides into the cytoplasm [30]. However, in the absence of all enzymes recognized to catalyze PGN recycling in *E. coli* the further fate of PGN turnover fragments remained elusive in *Chlamydiaceae*.

In this study, we aimed to gain mechanistic insight into recycling of PGN and found an NlpC/P60 ortholog protein to be encoded in *C. trachomatis*. This enzyme belongs to a primordial family of cell wall related peptidases that can be found throughout all kingdoms of life. Here, we show that *C. trachomatis* NlpC/P60 functions in recycling of PGN ring turnover products and specifically hydrolyzes PGN-derived peptides. We identify the cysteine-dependent enzyme as a promising target for antichlamydial intervention as it catalyzes a critical step in PGN recycling operating in the processes of energy recovery, cell division, and immune evasion.

## Results

### *C. trachomatis* harbors the homolog of a *Bacillus* NlpC/P60 protein

Searching the genome of *C. trachomatis* we found the pathogen to encode an NlpC/P60 protein, that shows 29% sequence identity with *Bacillus cereus* YkfC (YkfC_Bce_) and is hereinafter referred to as YkfC_Ctr_ (CT127). NlpC/P60 enzymes, including YkfC_Bce_, are cysteine peptidases that have a catalytic triad comprising Cys, His, and a polar residue [31]. Besides the C-terminal NlpC/P60 domain, YkfC_Bce_ contains two SH3b domains. The active site is located at the SH3b1-NlpC/P60 interface, whereas the SH3b2 domain is distal to the active site. Residues from the SH3b1 and NlpC/P60 domain form the so-called S2-S1 binding site cavity that determines the enzyme’s substrate specificity [31]. Our *in silico* sequence and structure analyses for YkfC_Ctr_ revealed that all catalytic residues and the S2-S1 binding cavity are conserved (Supplementary Fig. 1 and 12). Phyre^2^ predicts that these active site residues also form a catalytic center closely resembling the *B. cereus* YkfC one (Fig. 1a). The chlamydial homolog lacks membrane-spanning or lipoprotein-associated domains and a typical signal sequence for transport across the cytoplasmic membrane, suggesting a cytosolic localization of the enzyme. This may be supported by the identification of oligopeptide transporter system OppABCDF in *C. trachomatis* [30] which potentially transports PGN degradation-derived peptides into the chlamydial cell cytoplasm for further recycling as described for free-living bacteria.

**Figure 1.**
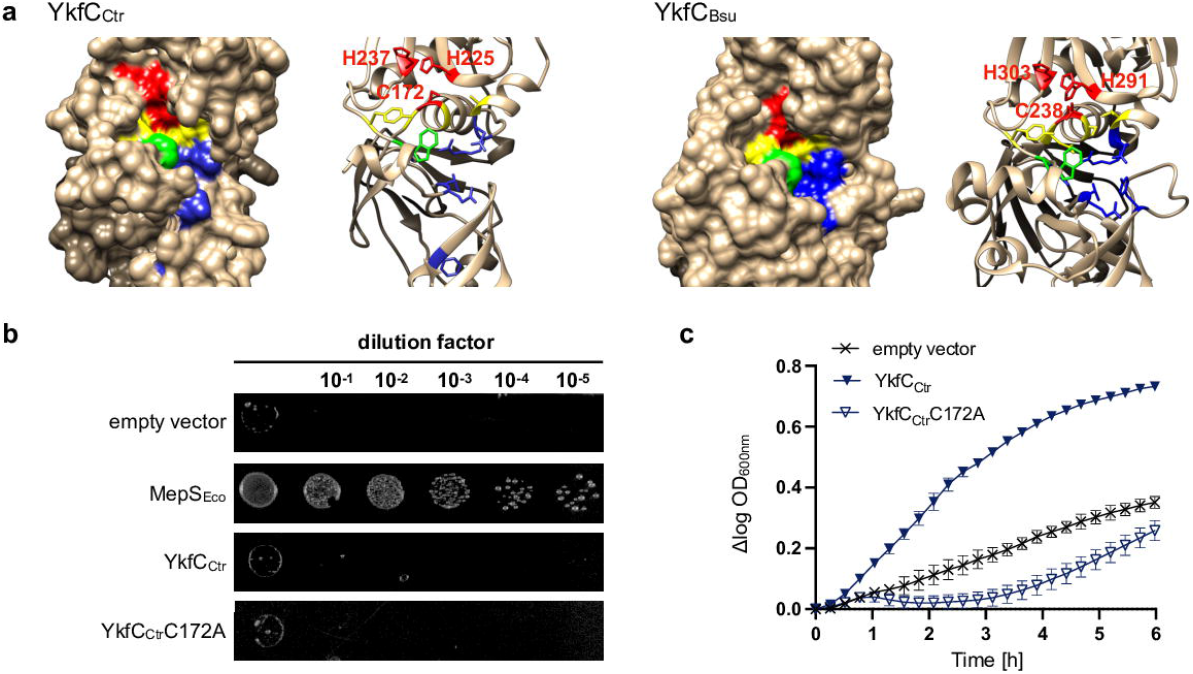
*In silico* analysis and activity of chlamydial YkfC in *E. coli*.□. **(a)** Comparison of the 3D *in silico* models of the active site from YkfC_Ctr_ and YkfC_Bce_ as characterized by Xu *et al*., 2010 [31]. The Cys-His-His catalytic triad is shown in red, residues contributing to the S1 and S2 sites essential for substrate recognition in YkfC_Bce_ are highlighted in yellow and blue, respectively. Residues contributing to both sites are highlighted in green. Complementation of *E. coli* Δ*mepS* and *E. coli* Δ*dapD*Δ*mpl* double mutant by chlamydial YkfC. **(b)** Spot plate assay of *E. coli* carrying a deletion for the gene encoding periplasmic NlpC/P60 endopeptidase MepS. The mutant was transformed with different pBAD24 constructs and grown at 42 °C on low osmolarity medium containing 0.04 % L-arabinose. Complementation of the osmo-and temperature-sensitive Δ*mepS* could only be observed for pBAD24 expressing *E. coli mepS*, but neither for *ykfC*_*Ctr*_ nor its active site mutant *ykfC*_*Ctr*_ (C172A). An empty vector control served as a negative control. **(c)** Growth effect in the *E. coli* Δ*dapD*Δ*mpl* double mutant containing chlamydial *ykfC* constructs. Cells were grown in nutrient-rich medium containing 40 µM PGN tripeptide and protein production was induced with 0.4 % L-arabinose. Cytoplasmic production of wild type YkfC_Ctr_, but not its active site mutant, restored growth of the *E. coli* double mutant, which is auxotrophic for mDAP and unable to utilize PGN tripeptide under nutrient rich conditions. Error bars indicate ± s.d. (n = 3).

### YkfC from *C. trachomatis* functions as a PGN tripeptide peptidase in *E. coli*

A number of the so far characterized cysteine-dependent NlpC/P60 proteins from free-living bacteria act as cell wall endopeptidases or peptide recycling enzymes [32]. To address these two possibilities for the *C. trachomatis* homolog, we tested if plasmid encoded YkfC_Ctr_ was able to restore growth in two different *E. coli* surrogate systems. First, we investigated whether YkfC_Ctr_ functions as a cell wall endopeptidase that cleaves cross-bridges between glycan chains within PGN as has been shown for other members of the NlpC/P60 family [31,32]. In *E. coli*, NlpC/P60 protein MepS (formerly Spr) is a DD-endopeptidase which specifically cleaves D-Ala-mDAP cross-links within PGN, thereby allowing incorporation of new glycan strands during cell growth [33]□. *E. coli* mutants lacking MepS show a thermosensitive growth defect at low osmolarity [34]. We took advantage of this phenotype to test for possible PGN endopeptidase activity of YkfC_Ctr_. Expression of neither wild type YkfC_Ctr_ nor the active site mutant YkfC_Ctr_C172A, in which the conserved Cys172 residue within the Cys-His-His triad was replaced by Ala, restored growth of an *E. coli* MepS mutant strain when grown at a non-permissive temperature and low osmolarity (Fig. 1b). These results make a PGN endopeptidase activity unlikely for YkfC_Ctr_ and suggest a role in recycling of PGN-derived peptides.

To test for recycling activity of the chlamydial NlpC/P60 protein we used an *E. coli* Δ*dapD*Δ*mpl* double mutant strain that reports on degradation of the PGN-derived tripeptide L-Ala-γ-D-Glu-mDAP through specific cleavage of the non-canonical γ-D-Glu-mDAP bond which is absent in proteins [35]. This strain is auxotrophic for Mdap and deficient in utilizing the PGN tripeptide. Auxotrophy for mDAP was achieved by deletion of *dapD*, which codes for a critical enzyme within the mDAP/L-Lys synthesis pathway. Direct incorporation of the tripeptide into newly synthesized PGN precursors was halted by deleting the gene *mpl* which encodes the ligase linking the tripeptide to the activated *N*-acetylamino sugar UDP-MurNAc, forming precursor UDP-MurNAc-L-Ala-γ-D-Glu-mDAP. We hindered the Δ*dapD*Δ*mpl* double mutant from hydrolyzing tripeptide through its peptidase MpaA that releases mDAP from the peptide by growing the cells under nutrient rich growth conditions in our test system. Under these conditions, the *E. coli* Δ*dapD*Δ*mpl* double mutant requires supplementation with mDAP for vital growth as expression of *mpaA* is diminished by the transcriptional regulator PgrR, affecting *E. coli* in hydrolyzing PGN tripeptide and thus in using it as a source of mDAP [35].

When incubated under nutrient rich growth conditions, the double mutant was severely impaired in using PGN tripeptide resulting in defective growth (Fig. 1c, empty vector control). The expression of YkfC_Ctr_ restored growth, suggesting that YkfC_Ctr_ functions as a PGN tripeptide peptidase in *E. coli* and released sufficient mDAP from the peptide to promote vital growth (Fig. 1c, Supplementary Fig. 2). To further investigate whether the observed growth promotion was connected to the catalytic activity of YkfC_Ctr_, we expressed the active site mutant YkfC_Ctr_C172A under the same conditions. The active site mutant YkfC_Ctr_C172A was unable to restore growth in the surrogate system, indicating that the observed growth effect depended on the presence of catalytically active YkfC_Ctr_ (Fig. 1c).

In sum, our observations suggest that YkfC_Ctr_ is a functional γ-D-Glu-mDAP peptidase when expressed in *E. coli* and that it operates in recycling of tripeptides released during PGN degradation.

### YkfC_Ctr_ is active *in vitro*

To further characterize the chlamydial NlpC/P60 homolog at a molecular level we proceeded with biochemical analyses using the heterologously overproduced and purified enzyme (Supplementary Fig. 3a). Consistent with our results from the surrogate test system, the purified enzyme used PGN tripeptide as a substrate. Our thin-layer chromatography (TLC) analysis showed that wildtype YkfC_Ctr_ was able to release mDAP from the PGN tripeptide L-Ala-γ-D-Glu-mDAP by hydrolyzing the γ-D-Glu-mDAP bond (Fig. 2a,b). This non-protein bond, which is found in Gram-negative PGN and its degradation products, binds the lateral (γ)-carboxyl group of a D-amino acid (D-Glu) to the α-amino group of the non-proteinogenic amino acid mDAP [27]. The enzyme’s activity was time-dependent (Supplementary Fig. 3b) and showed a pH optimum of 7.5-8.5 (Fig. 2c). Consistent with our observation that the expression of YkfC_Ctr_C172A failed to rescue growth in the *E. coli* Δ*dapD*Δ*mpl* surrogate system, the purified protein variant was inactive against L-Ala-γ-D-Glu-mDAP (Fig. 2b). PGN recycling needs to be tightly controlled and product inhibition could provide a negative feedback. To investigate possible product inhibition, we also tested if mDAP inhibits the enzyme in concentrations comparable to the substrate concentration. mDAP did not show any inhibitory effect on the enzyme, suggesting negative feedback through this reaction product does not occur (Supplementary Fig. 3c). As we could not completely rule out that the heterologous YkfC_Ctr_ produced in *E. coli* was contaminated by the zinc dependent host peptidase MpaA which is also capable of releasing mDAP from the PGN tripeptide, we investigated whether the metal chelator EDTA had any effect in our assay. In accordance with our active site mutant experiments that did not reveal background activity, we could not detect any inhibition of substrate turnover under these conditions, further attesting for high purity and the absence of interfering host peptidases in our protein purification (Supplementary Fig. 3d).

**Figure 2.**
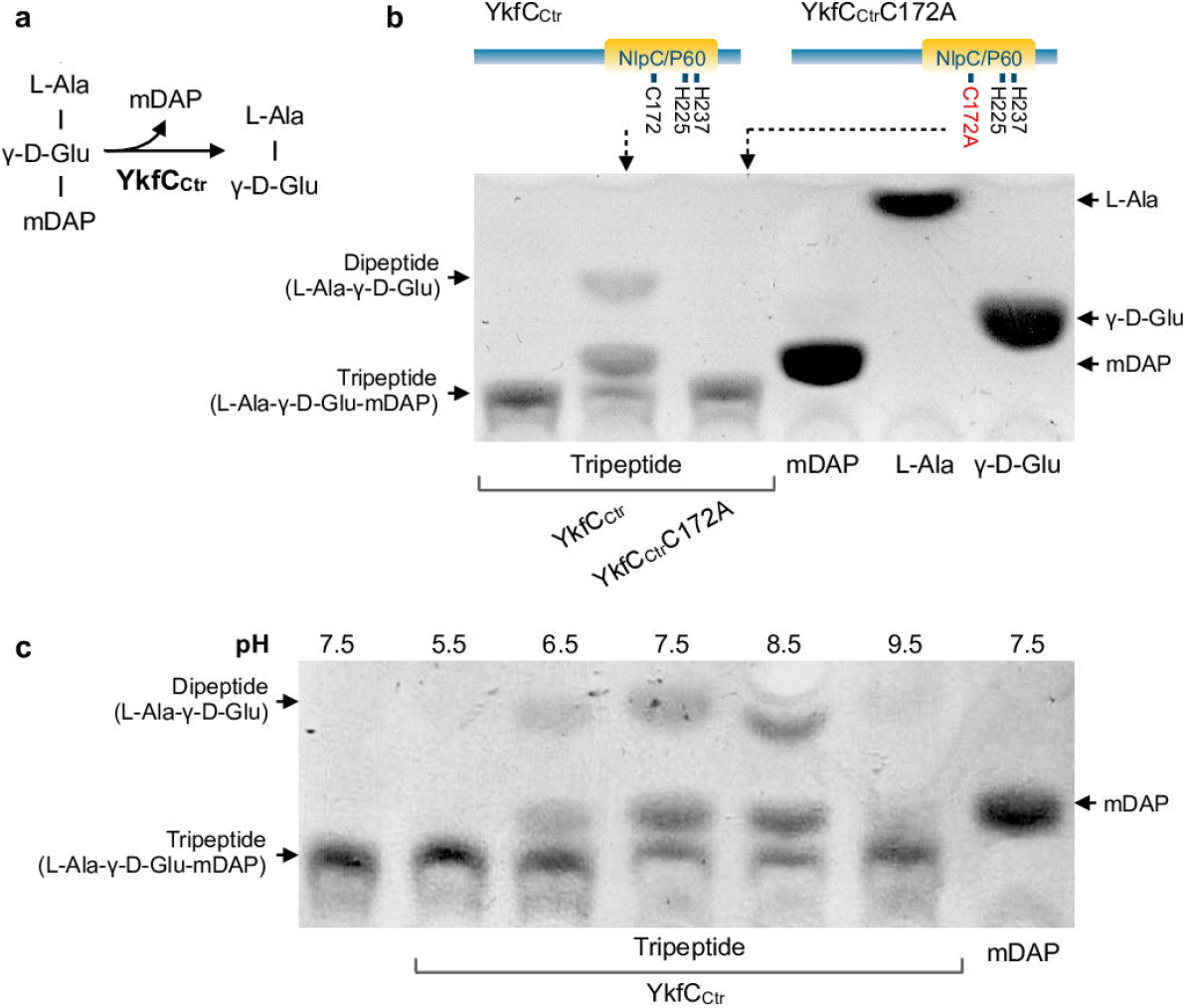
*In vitro* activity of YkfC_*Ctr*_. **(a)** Schematic representation of the proposed peptidase activity of YkfC_Ctr_ releasing mDAP from L-Ala-γ-D-Glu-mDAP tripeptide. **(b)** *In vitro* peptidase activity was analyzed by TLC using L-Ala-γ-D-Glu-mDAP tripeptide as a substrate. YkfC_Ctr_, but not its active site mutant YkfC_Ctr_C172A, released mDAP from the tripeptide as shown by comparing migration of the reaction product and an mDAP standard. **(c)** pH dependence of YkfC_Ctr_ activity. The chlamydial YkfC homolog showed highest activity between pH 7.5 and 8.5 and was inactive at acidic pH 5.5.

Our results identified Ykfc_Ctr_ as a peptidase that cleaves mDAP from PGN tripeptide thus corroborating its role in recycling PGN-derived degradation products.

### YkfC_Ctr_ is specific for PGN-derived peptides

To substantiate that YkfC_Ctr_ is dedicated to recycling of PGN degradation products, we screened different PGN-derived peptides and intermediates from the *de novo* PGN precursor biosynthesis pathway as potential substrates (Fig. 3). Testing PGN-derived peptides, we found the enzyme to be active not only against the tripeptide L-Ala-γ-D-Glu-mDAP, but also against the tetrapeptide L-Ala-γ-D-Glu-mDAP-D-Ala, another typical degradation product released by PGN amidases such as chlamydial AmiA [20]. As evidenced by our MS analyses, cleavage in the L-Ala-γ-D-Glu-mDAP tripeptide and the L-Ala-γ-D-Glu-mDAP-D-Ala tetrapeptide only occurred between D-Glu and mDAP and neither between L-Ala and D-Glu nor between mDAP and D-Ala, when the tetrapeptide was used as a substrate. These data further supported that the enzyme specifically cleaves γ-D-Glu-mDAP bonds. Modification of the tripeptide through mDAP amidation at the free carboxyl group as in *Bacillus* PGN did not affect the enzyme (Supplementary Fig. 4). However, removal of L-Ala in position 1, resulting in dipeptide γ-D-Glu-mDAP, abolished activity (Fig. 3a). NlpC/P60 recycling enzymes from free-living bacteria were described to depend on a free L-Ala terminus for proper substrate recognition [32]. To test whether this requirement applies to chlamydial YkfC, we next investigated PGN intermediates where L-Ala in position 1 of the peptide is linked to sugar moieties. Indeed, YkfC_Ctr_ showed no activity on the final PGN building block lipid II, the nucleotide-activated precursor UDP-MurNAc-tripeptide or the PGN recycling intermediate MurNAc-tripeptide (Fig. 3a,c). To further elaborate on the enzyme’s specificity, we additionally tested a set of 10 (non)-PGN peptides which did not fulfill the above determined requirement combining the presence of a γ-D-Glu-mDAP bond and a free L-Ala terminus. None of the peptides was used as a substrate by YkfC_Ctr_ (Supplementary Fig. 5). YkfC_Ctr_ had also no activity on high molecular weigth PGN in remazol dye release assays, which is consistent with the *E. coli* complementation experiments and further supports a function in recycling PGN-derived peptides in the cytoplasm (Fig. 3d).

**Figure 3.**
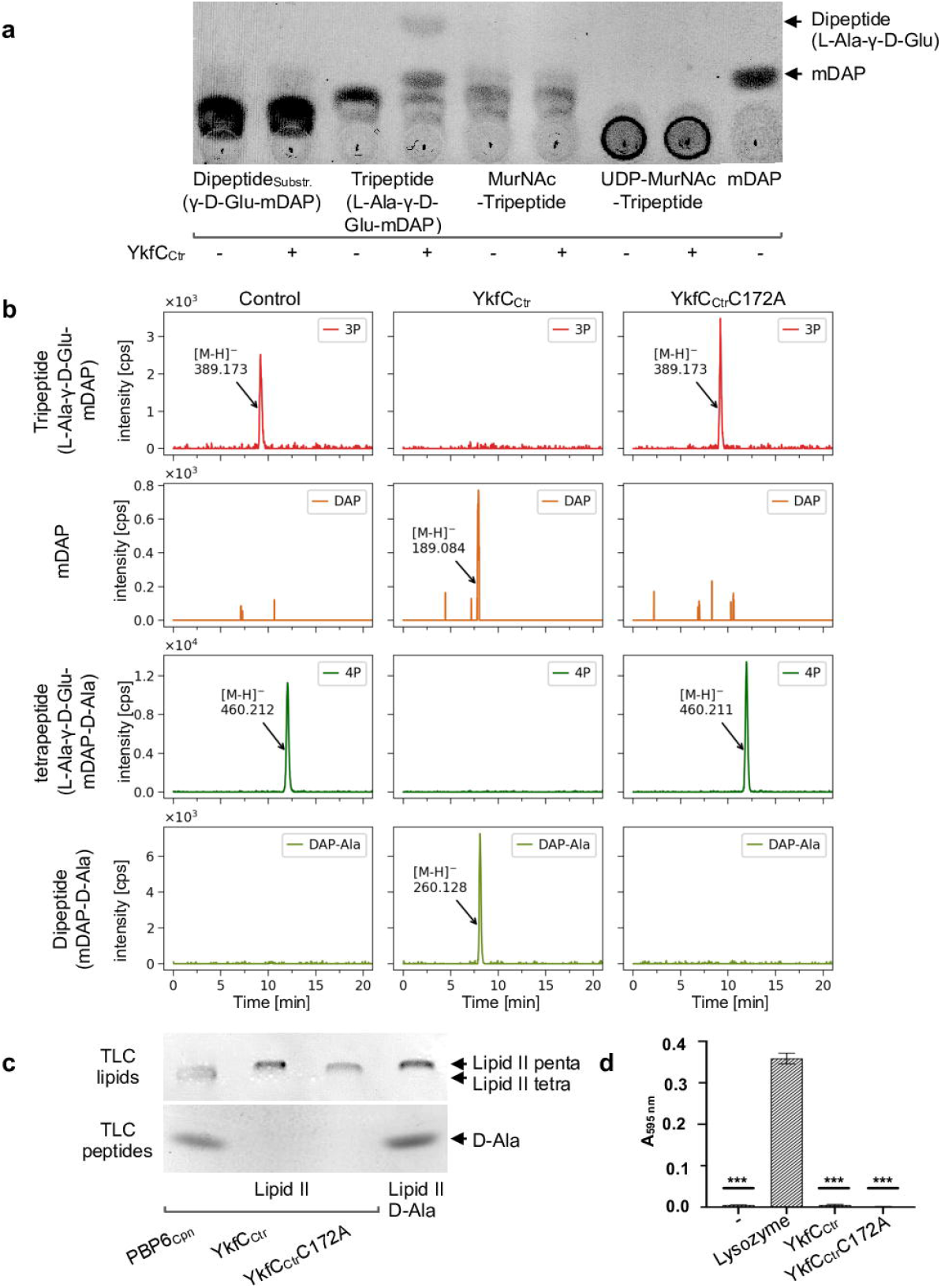
Substrate specificity of YkfC_Ctr_. YkfC_Ctr_ hydrolyzes the bond between D-Glu and mDAP in the PGN tri-and tetrapeptide. **(a)** *In vitro* activity of YkfC_Ctr_ on soluble precursors and recycling products. TLC analysis shows that YkfC_Ctr_ uses PGN-derived tripeptide as a substrate but neither the lipid II precursor UDP-MurNAc-tripeptide nor the PGN recycling intermediate MurNAc-tripeptide. PGN tripeptide is the minimal substrate as the dipeptide γ-D-Glu-mDAP was not hydrolyzed under our test conditions. **(b)** LC-MS analysis of YkfC_Ctr_ activity on CwlC amidase derived peptides from *E. coli* PGN. YkfC_Ctr_, but not its active-site mutant, cleaves the tri-and tetrapeptide as shown by the disappearance of these substrates and the production of mDAP and mDAP-D-Ala products, respectively. Data are presented as extracted ion chromatograms applying the theoretical masses of tripeptide (3P, [M−H]^−^ = 389.167), tetrapeptide (4P, [M−H]^−^ = 460.204), DAP ([M−H]^−^ = 189.087) and DAP-Ala ([M−H]^−^ = 260.124) within an error range of ± 0.02. **(c)** *In vitro* activity of YkfC_Ctr_ on the membrane bound PGN precursor lipid II. Neither YkfC_Ctr_ nor its active site mutant YkfC_Ctr_C172A exhibit detectable activity on PGN precursor lipid II as shown by the unaltered migration distance on the lipid TLC and the absence of amino acid or peptide reaction products on the peptide TLC. DD-carboxypeptidase PBP6_Cpn_ releases the terminal D-Ala from lipid II and was included as a positive control to show that the employed TLC method is suitable to detect changes within the structure of lipid II (lipid TLC) or the release of amino acids and peptides from lipid II (peptide TLC). **(d)** Dye-release assay and photometric analysis of the reaction products. YkfC_Ctr_ and its active-site mutant YkfC_Ctr_C172A did not exhibit hydrolytic activity when incubated with remazol-brilliant-blue labelled PGN. Lysozyme served as a positive control releasing the labelled muropeptides into the supernatant. Unpaired t-test revealed statistical significance in comparison with the lysozyme control, two-tailed \*\*\**P*-value ≤0.0001. Error bars indicate ± s.d. (n=3). (lipid II penta: undecaprenyl-pyrophosphoryl-MurNAc-(GlcNAc)-L-Ala-γ-D-Glu-mDAP-D-Ala-D-Ala; lipid II tetra: undecaprenyl-pyrophosphoryl-MurNAc-(GlcNAc)-L-Ala-γ-D-Glu-mDAP-D-Ala).

Taken together, we found evidence that YkfC from *C. trachomatis* is tailored for recycling of PGN peptides that are released from the PGN glycan chains by amidases. Provided that the peptides meet the requirement of both a γ-D-Glu-mDAP bond and a free L-Ala terminus, the enzyme appeared to show a rather relaxed substrate spectrum as attested by use of an amidated mDAP variant. Precluding hydrolysis of PGN biosynthesis intermediates such as lipid II and its precursor UDP-MurNAc-tripeptide, where the N-terminus is occupied by an *N*-acetylmuramyl residue, guarantees that cytosolic YkfC_Ctr_ does not interfere with intracellular lipid II biosynthesis as otherwise, the tightly orchestrated processes of PGN ring biosynthesis and cell division would collapse.

### YkfC_Ctr_ is inhibited by the alkylating agent chloroacetone and the peptidomimetic epoxide E-64

Both our *in vivo* experiments in the *E. coli* surrogate system and biochemical analyses with the purified protein pointed to a crucial role of the conserved cysteine C172 within the NlpC/P60 active site. We used a set of cysteine protease inhibitors as chemical tools to further explore the role of cysteine C172 in the catalytic activity of YkfC_Ctr_. Chlamydial proteins typically contain a high number of cysteines which are otherwise relatively rare amino acids [18,36]. Besides the cysteine present in the predicted active site, YkfC_Ctr_ contains 7 additional cysteines of unknown redox state that could be targeted by alkylating compounds which bind to all accessible sulfhydryl-groups. For this reason, we tested inhibitor concentrations for these compounds in relation to the total number of all theoretically available cysteine residues. For the cysteine protease inhibitor E-64 this adaptation was not necessary as the compound selectively binds to the active site of cysteine proteases [37]. Among the two tested alkylating compounds chloroacetone and iodoacetamide, only chloroacetone inhibited the enzyme at a molar inhibitor:protein ratio of 8:1 (Fig. 4a). Iodoacetamide did not inhibit YkfC_Ctr_ even at molar ratios of 80:1 (Fig. 4b). The same specificity has been described for the YkfC homolog from *Bacteroides ovatus* [32]□ and the lack of inhibition by iodoacetamide is likely attributed to the significantly lower electronegativity of the iodine compared to the chlorine atom in chloroacetone. The irreversible cysteine protease specific inhibitor E-64 abolished YkfC_Ctr_ activity at a molar inhibitor:protein ratio of 10:1 (Fig. 4d). These data suggest an essential role of the active site cysteine. Future crystallization studies might reveal the overall architecture of the enzyme’s active site and provide detailed information on substrate and inhibitor specificity and recognition.

**Figure 4:**
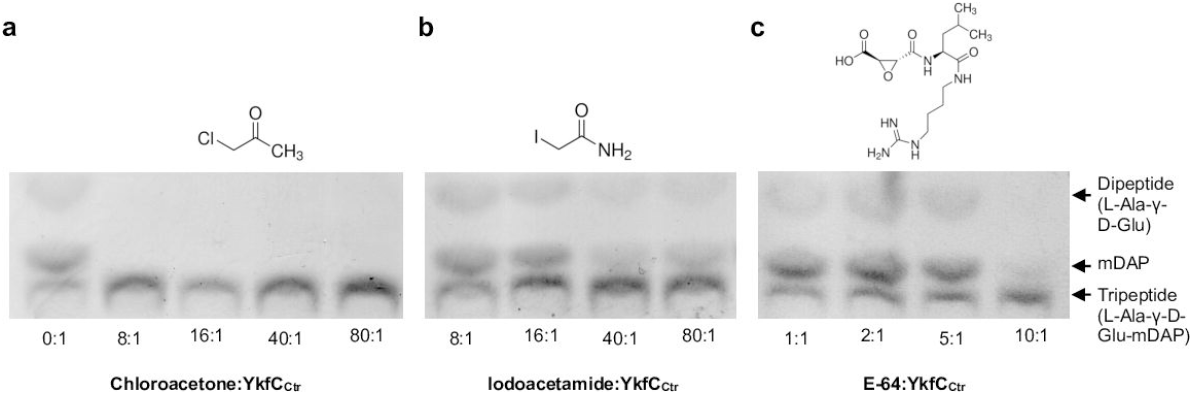
Effect of different inhibitors on the enzymatic activity of YkfC_Ctr_. TLC analysis of the inhibiting effect of chloroacetone, iodoacetamide, and E-64 on YkfC_Ctr_ activity. **(a)** The alkylating chemical chloroacetone proved to be the most potent inhibitor as shown by the absence of the reaction product mDAP at a molar inhibitor:protein ratio of 8:1. YkfC_Ctr_ retained activity at lower molar ratios (Supplementary Fig. 6). **(b)** The second alkylating compound iodoacetamide did not show a significant inhibitory effect on YkfC_Ctr_ activity. **(c)** The cysteine protease inhibitor E-64 carrying a *trans*-epoxysuccinic acid group coupled to a dipeptide was able to inhibit YkfC_Ctr_ at a molar ratio of 10:1.

### Chloroacetone and E-64d have anti-chlamydial activity in cell culture infection experiments

Our *in vitro* results on an inhibitory effect of chloroacetone and natural compound E-64 on purified YkfC_Ctr_ prompted us to test for *in vivo* activity of these cysteine protease inhibitors against *C. trachomatis* in cell culture experiments. Experiments were performed in the range of inhibitor concentrations and treatment durations that did not exert cytotoxicity towards the HEp-2 host cells in resazurin-based viability assays (Supplementary Fig. 7). We propagated *C. trachomatis* D in human HEp-2 cells, added inhibitors at 6, 10 and 24 hpi and analyzed chlamydial infection at 30 hpi using immunofluorescence microscopy (leading to treatment durations of 24, 20 and 6 h, respectively, Fig. 5). E-64 did not show antichlamydial activity up to the highest concentration tested (256 μg/mL) irrespective of treatment duration, probably due to a poor penetration of the lead compound into chlamydial cells. Also cytotoxicity could not be observed for E-64 up to the highest concentration (256 μg/mL) and treatment duration (24h) tested. The more membrane-permeable variant E-64d [38] had antichlamydial activity, reflected by a minimal inhibitory concentration (MIC) of 256 μg/mL after treatment for 6 h (corresponding to an addition of the compound at 24 hpi). Under this treatment regimen, the compound did not exert cytotoxicity towards the host cells but had a detrimental effect on chlamydial cells culmulating in the disruption of chlamydial inclusions and cells -possibly due to its concerted interference with several cellular processes. Analyses with extended E-64d treatment durations were limited by first cytotoxic effects of the variant at a concentration of 256 μg/mL. For E-64d treatment durations of 20 and 24 hpi (corresponding to an addition of the compound at 10 and 6 hpi), we observed no significant reduction in the inclusion number at 128 μg/mL, the highest concentration without cytotoxicity after these prolonged treatment durations. Chloroacetone had the most pronounced antichlamydial activity resulting in a MIC of 4 μg/mL for all tested treatment durations and without detectable impairment of host cell viability in the range of MIC (Supplementary Fig. 7).

In a complementary approach, we employed a dCas12 CRISPR interference-based inducible knockdown system [39]. We verified knockdown of *ykfC*_*Ctr*_ in *C. trachomatis* L2 by measuring the transcriptional levels of *ykfC*_*Ctr*_ in comparison to *euo* and *omcB*, as controls for developmental cycle progression [40] (Supplementary Fig. 8a). We also assessed genomic DNA levels and did not observe significant differences after induction of dCas12 expression (Supplementary Fig. 8b). Expression of the dCas12 in the non-targeting background resulted in an approximately 50% reduction in infection forming units, suggesting there is a metabolic burden to overexpressing dCas12 that does not impact replication rate (Supplementary Fig. 8b,c). Indirect immunofluorescence assay (IFA) analysis did not reveal a meaningful change in inclusion and chlamydial morphology (Supplementary Fig. 8d,e). Nonetheless, after knocking down *ykfC*_*Ctr*_ transcripts, there was a further, statistically significant 50% reduction in production of infection forming units in the *ykfC* knockdown strain as compared with the non-targeting control (Supplementary Fig. 8c). The less natural environment in the permissive mouse fibroblast McCoy cells and residual expression and enzymatic activity of YkfC_Ctr_ might have contributed to the less intense effects in the *ykfC* knockdown experiments.

**Figure 5.**
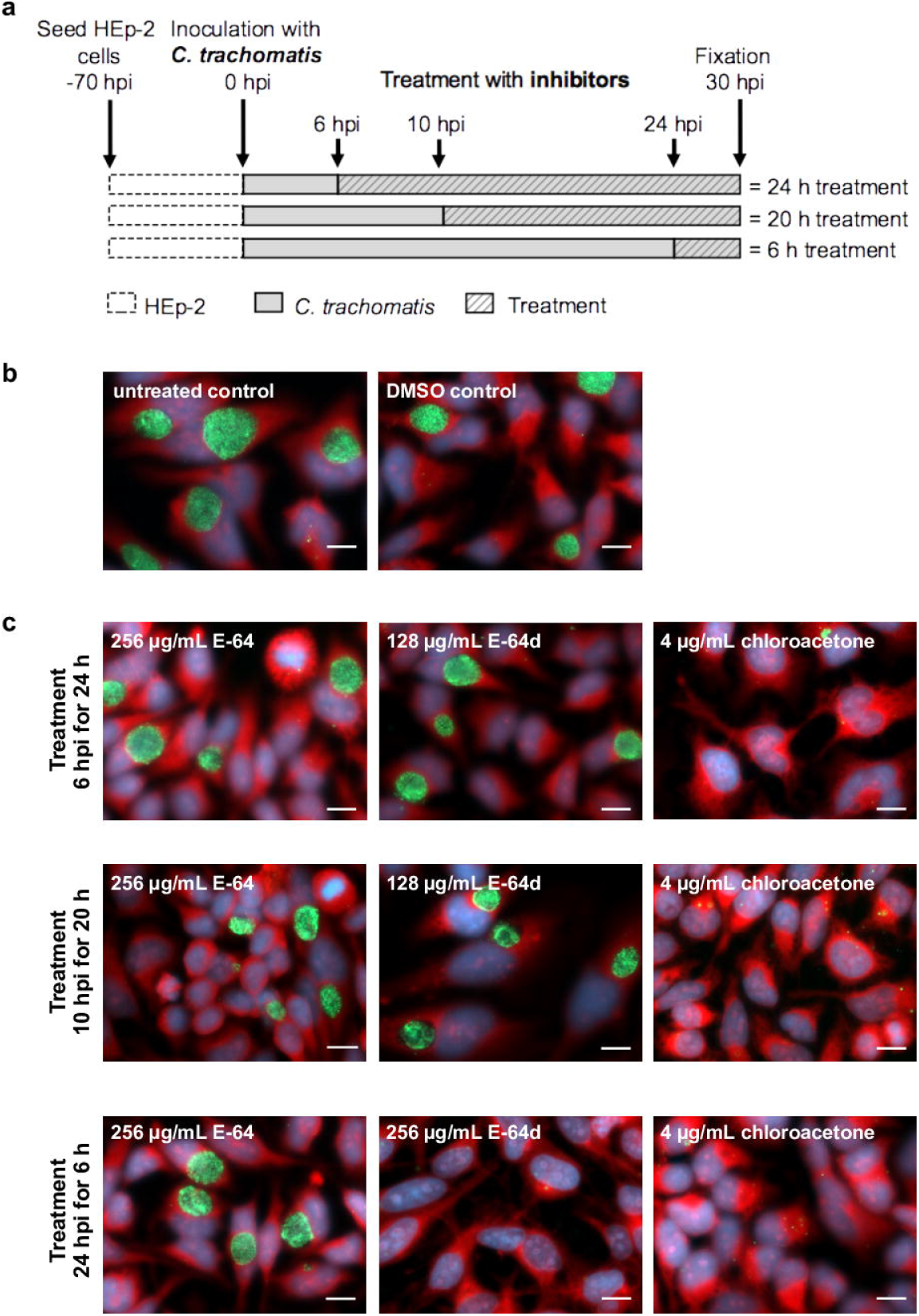
Determination of minimal inhibitory concentrations of YkfC inhibitors towards *C. trachomatis*. Overview of the inhibitor susceptibility analyses **(a)**. HEp-2 cells were infected with *C. trachomatis* serovar D strain UW-3/CX, E-64, its variant E-64d, and chloroacetone were added at 6, 10 and 24 hpi and the antichlamydial activity was analyzed at 30 hpi (resulting in total treatment durations of 24, 20 and 6 h, respectively). Experiments were done in triplicate and analyzed in the range of concentrations and treatment durations that had no cytotoxicity towards the HEp-2 host cells (Supplementary Fig. 7). Exemplarily pictures of untreated and DMSO-treated **(b)** as well as inhibitor-treated cells at the minimal inhibitory concentration (MIC) or the highest concentration tested **(c)** were included in the figure. The MIC was the lowest concentration that reduced the number of chlamydial inclusions by more than 90% as defined previously [55]. E-64 had neither antichlamydial nor cytotoxic activity up to the highest concentration and treatment duration tested (256 μg/mL for 24 h). In contrast, E-64d treatment for 6h severely impaired chlamydial infection without cytotoxicity resulting in a MIC of 256 μg/mL. Extended E-64d treatment with 256 μg/mL for 20 and 24 h (corresponding to an addition at 6 and 10 hpi) lead to reduced host cell viability. Treatment with 128 μg/mL, the highest concentration without cytotoxicity under these treatment durations, had only a minor non-significant antichlamydial effect. With an MIC of 4 μg/mL determined for all tested treatment durations, chloroacetone had the highest antichlamydial activity without cytotoxicity towards host cells in the range of MIC.

All in all, we used YkfC_Ctr_ inhibitors as tools in *in vivo* cell culture studies and provided further evidence that the NlpC/P60 protein is important for chlamydial biology. In particular, the inhibitory effect of the more membrane-permeable variant of natural lead compound E-64, although less pronounced compared to chloroacetone, may open the way for further studies on YkfC_Ctr_ as a target for antichlamydial drug development.

### Localization of YkfC in *C. trachomatis* cells

To further explore the biological function of YkfC_Ctr_ we performed localization analyses in *C. trachomatis* (Fig. 6). To this end, we made an anhydrotetracycline (aTc)-inducible construct encoding YkfC_Ctr_ tagged with six histidines at the C-terminus (YkfC_Ctr__6xH). This construct was transformed into *C. trachomatis* serovar L2 lacking its endogenous plasmid (-pL2). Using indirect immunofluorescence assays (IFA), we detected YkfC_Ctr__6xH as puncta at the periphery of non-dividing *C. trachomatis* cells (Fig. 6b). We did not observe any gross morphological effects of YkfC_Ctr__6xH overexpression on inclusion size or bacterial numbers, as confirmed by inclusion forming unit assays where the YkfC_Ctr__6xH expressing strain generated 96 ± 20% of infection forming units compared to the uninduced control (Supplementary Fig. 9). These data indicate that Ykf_Ctr__6xH overexpression does not impact chlamydial developmental cycle progression.

**Figure 6.**
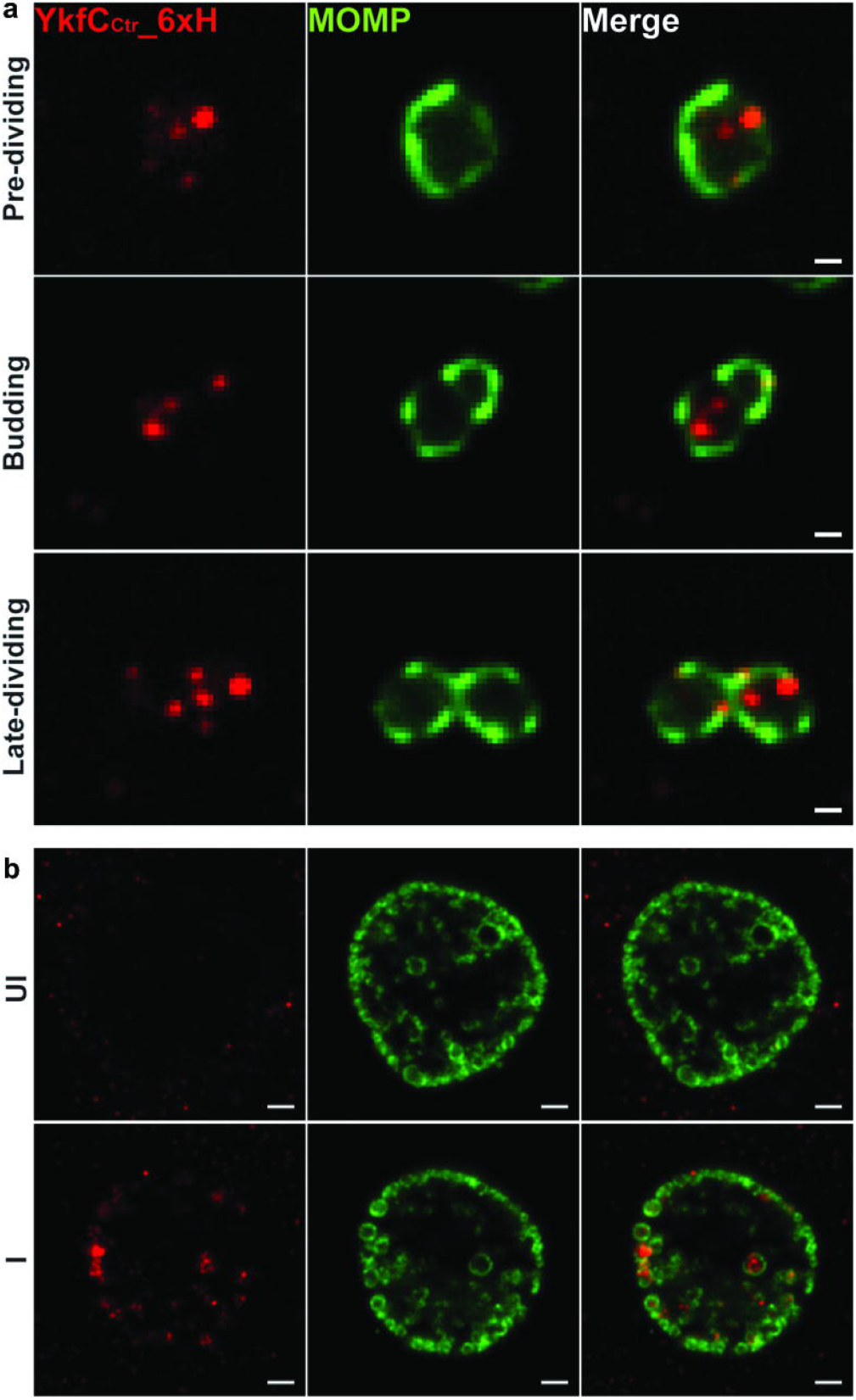
Localization of YkfC_Ctr__6xH in *C. trachomatis*. McCoy cells were infect-ed with the transformant encoding YkfC_Ctr__6xH. YkfC_Ctr__6xH expression was induced at 4 hpi with 10 nM aTc. At 10.5 **(a)** or 24 hpi **(b)**, the infected cells were fixed with aldehydes to preserve the budding morphology of dividing cell (10.5 hpi) or 100% MeOH (24 hpi) and stained with anti-6xH (red) and anti-MOMP (green) antibodies. The images were acquired on a Zeiss AxioImager.Z2 equipped with an Apotome2 using a 100X objective. Scale bar: 0.5 μm **(a)** or 2 μm **(b)**. YkfC_Ctr__6xH showed as puncta at the periphery of cells. As exemplarily shown in **(a)**, assessment of dividing cells revelaed that 90% of cells in the early stages of cell division (19 out of 21 ana-lyzed cells in the pre-dividing and following budding stage) did not feature YkfC_Ctr__6xH signals at the (future) site of division (i.e. at the MOMP-enriched site in pre-dividing cells or at the site of daughter cell formation in the budding stage). In contrast, 60% of cells in the late dividing stage (15 out of 25 analyzed late diving cells) had YkfC_Ctr__6xH puncta near the septum. UI: uninfected; I: infected.

When analyzing dividing cells, we found evidence for an overall enrichment of the Ykf_Ctr__6xH signal near the septum as cell division progresses (Fig 6a). Only 10% of cells in the early stages of cell division featured Ykf_Ctr__6xH signals proximal to the site of division (i.e. at the MOMP (major outer membrane protein)-enriched site which determines the future site of daughter cell formation [22,41] in the pre-dividing stage or at the site of polarized asymmetric daughter cell formation in the following budding stage). In contrast, Ykf_Ctr__6xH puncta near the septum could be detected in 60% of cells in the late cell-division stage. These observations suggest that YkfC_Ctr_ might be recruited adjacent to the cell division plane later in the process, consistent with the function of an enzyme which is involved in recycling of PG material, a process that is likely coordinated with constriction and degradation of the transient PG ring during separation of daughter cells.

### NOD1 activation by *C. trachomatis* is increased under YkfC inhibitor treatment

PGN-derived peptides are known to activate innate immunity factor NOD1 that binds to the γ-D-Glu-mDAP dipeptide as a minimal ligand [42]. We wanted to learn whether YkfC_Ctr_ is active *in vivo* and contributes to immune evasion through degrading the recognition motif of NOD1 in PGN-derived peptides. To this end, we performed NOD1 activation experiments with *C. trachomatis* infected human NOD1 reporter cells in the presence of an YkfC inhibitor. Indeed, NOD1 activation was significantly increased under treatment with the YkfC inhibiting compound chloroacetone as compared with the untreated infected vehicle control (Fig. 7). The increase of NOD1 activation was observed at concentrations (0.016 and 0.031 μg/mL) that did not impact on inclusion size (Supplementary Fig. 10a) and were below both the MIC of chloroacetone towards *C. trachomatis* (4 μg/mL) and cytotoxicity towards the infected reporter cell line (IC_50_ 15 μg/mL, Supplementary Fig. 10b). In control experiments with uninfected NOD1 reporter cells, chloroacetone did not modulate the level of NOD1 activation (Supplementary Fig. 10c). These results provide evidence that *C. trachomatis* YkfC degrades NOD1 stimulating PGN-derived peptides not only *in vitro* but also in infected host cells supporting a role in immune evasion.

**Figure 7.**
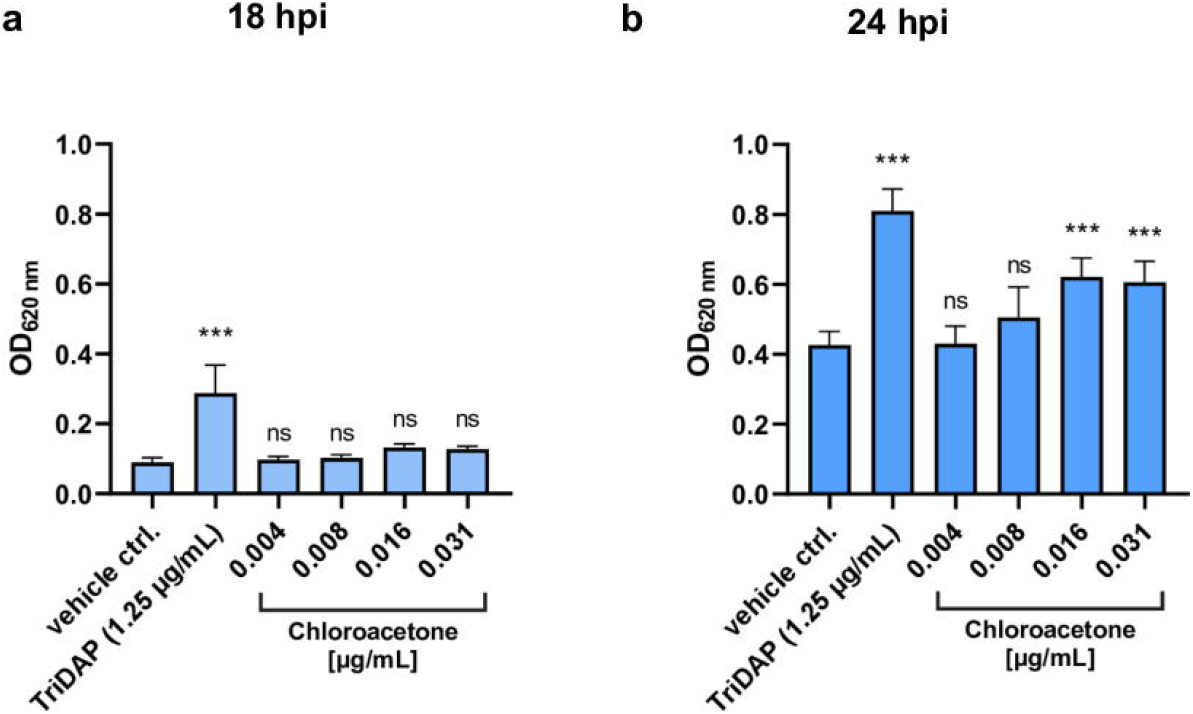
NOD1 signaling in *C. trachomatis* infected reporter cells is increased under YkfC inhibitor treatment. NOD1 reporter cells were infected with *C. trachomatis* D and used to analyze abundance of NOD1 activating PGN-derived material shed by the intracellular pathogen in the presence of YkfC_Ctr_ inhibitor chloroacetone. Infected NOD1 reporter cells were treated 6 hpi with chloroacetone and analyzed 18 hpi **(a)** and 24 hpi **(b)** resulting in treatment durations of 12 h and 18 h, respectively. An increase of NOD1 activation was observed at concentrations (0.016 and 0.031 μg/mL) below both cytotoxicity towards the reporter cell line (IC_50_ 15 μg/mL after 24 h treatment, Supplementary Fig. 10a) and the MIC of chloroacetone towards *C. trachomatis* (4 μg/mL, observed after treatment durations of 6, 20 and 24 hpi, Fig 5). Addition of TriDAP 6 hpi was used as a positive control for NOD1 activation. Error bars indicate ± s.d. (n=3). Ns: not significant. ***P-value ≤0.0001.

### NlpC/P60 proteins are widely conserved within the order *Chlamydiales*

Our discovery of a functional YkfC homolog in *C. trachomatis* prompted us to analyze the evolutionary conservation of NplC/P60 proteins among the *Chlamydiales*. Using sequences of YkfC from *C. trachomatis* and *B. cereus* as queries for BLAST alignments, we found genes coding for putative NlpC/P60 proteins in all members of the *Chlamydiaceae* and the genomically less reduced *Chlamydia*-related bacteria belonging to the *Simkaniaceae, Parachlamydiaceae*, and *Criblamydiaceae* families, but not within the family *Waddliaceae* [1,43]. To gain insight into the function of these putative NlpC/P60 proteins, we selected three candidates for initial analyses in our two *E. coli* surrogate systems reporting on possible PGN endopeptidase or peptide recycling activity.

Consistent with our results for *C. trachomatis* YkfC, neither the homolog from the other established human pathogen *C. pneumoniae* nor the putative NlpC/P60 proteins from the *Chlamydia*-related bacteria *Simkania negevensis* and *Estrella lausannensis* rescued growth in the endopeptidase MepS defective *E. coli* strain (Fig. 8a), making a role in periplasmic PGN turnover unlikely for all three candidates. The homolog from *C. pneumoniae* did not restore growth of the *E. coli* Δ*dapD*Δ*mpl* double mutant strain under nutrient-rich growth conditions (Fig. 8b). Both NlpC/P60 proteins from the *Chlamydia*-related organisms lack a typical signal sequence for transport across the cytoplasmic membrane and were capable of restoring growth in the *E. coli* surrogate test strain suggesting their ability to cleave PGN-derived tripeptide (Fig. 8b). The predicted structure of *S. negevensis* NlpC/P60 (NlpC/P60_Sne_) contrasts with all other chlamydial proteins. NlpC/P60_Sne_ lacks both SH3b domains including the S2 site residues (corresponding to Glu83, Thr84, and Tyr118 in YkfC_Bce_) important for interactions with L-Ala in position 1 of PGN peptides in other NlpC/P60 PGN peptidases. The structure predictions show highest matches with an NlpC/P60 PGN hydrolase from the protozoan *Trichomonas vaginalis* (Supplementary Fig. 12 and 13) [44]. These data indicate that our understanding of substrate binding in prokaryotic NlpC/P60 proteins is not yet fully established. Moreover, our results suggest that NlpC/P60 proteins are likely conserved as peptide recycling enzymes not only in *C. trachomatis* but also the less related, deeply-rooting *S. negevensis* and *E. lausannensis*.

**Figure 8.**
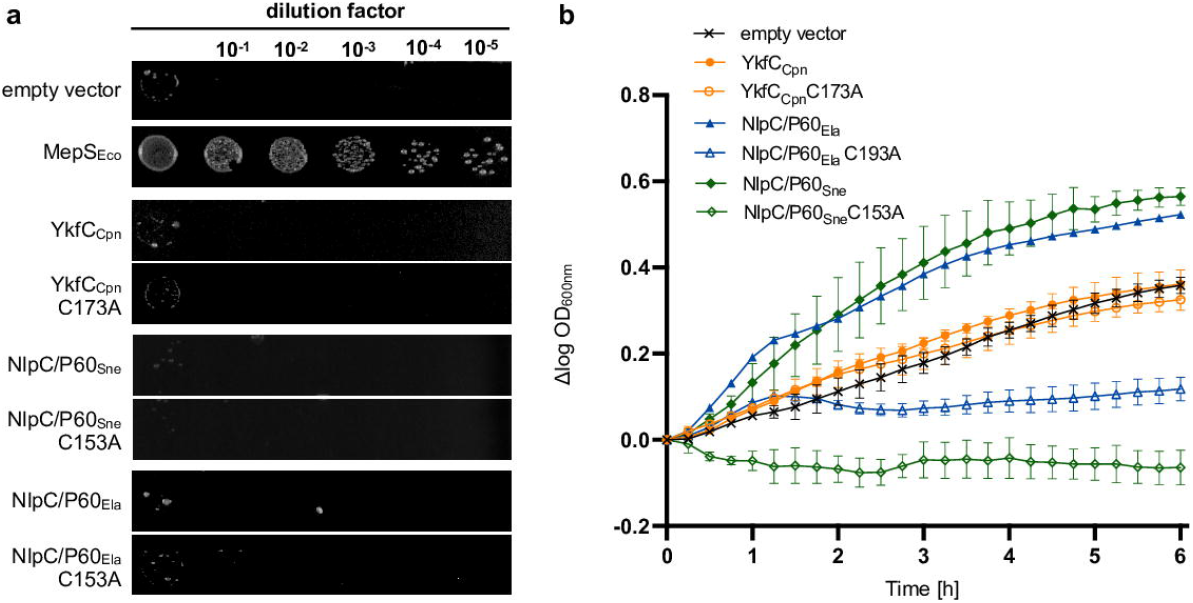
*In vivo* complementation of the *E. coli* Δ*dapD*Δ*mpl* and *E. coli* Δ*mepS* surrogate systems with different chlamydial NlpC/P60 homologs. **(a)** Spot plate assay of *E. coli* carrying a deletion for the gene encoding periplasmic NlpC/P60 endopeptidase MepS. The mutant was transformed with different pBAD24 constructs encoding for an NlpC/P60 homolog from *C. pneumonia, S. negevensis*or *E. lausannensis* and their cognate active-site mutants. Plates with low osmolarity media containing 0.04 % L-arabinose were incubated at 42 °C. Complementation of the osmo- and temperature-sensitive Δ*mepS* could only be observed for pBAD24 producing *E. coli* MepS, but not for any chlamydial NlpC/P60 homolog. An empty vector control served as a negative control. **(b)** Growth effect in the *E. coli* Δ*dapD*Δ*mpl* double mutant containing different chlamydial NlpC/P60 constructs. Cells were grown in nutrient-rich medium containing 40 µM PGN tripeptide and protein production was induced with 0.4 % arabinose. NlpC/P60 from *S. negevensis* and *E. lausannensis*, but not from *C. pneumoniae*, restored growth of the *E. coli* double mutant, which is auxotrophic for mDAP and unable to utilize PGN tripeptide under nutrient rich conditions. None of the active site mutants was able to complement the growth defect. Error bars indicate ± s.d. (n = 3).

To further explore the function of NlpC/P60 proteins in the biology of both *Chlamydia*-related genera, we analyzed gene expression and protein localization of of NlpC/P60 homologs in *S. negevensis* and *E. lausannensis* (Fig. 9a). Transcription of *nlpC/p60* could be detected in both species (Fig. 9a) and was normalized to the number of bacteria, as quantified by qPCR (Supplementary Fig. 14a). In *E. lausannensis* the expression level of *nlpC/p60* was similar to *mreB*, and both transcript levels peaked synchronously during cell divsion, as expected. To date, nothing is known about cell division in *S. negevensis*. As described above, our *E. coli* complementation experiments indicate that *S. negevensis* NlpC/P60 is capable of recycling PGN-derived peptides like the other tested proteins but the predicted structure contrasted with the other peptidases. In *S. negevensis, mreB* showed a less pronounced increase in expression during the replication phase as compared with *E. lausannensis*, and also the *nlpC/p60* gene was more constantly expressed throughout the developmental cycle. This might imply functional differences between NlpC/P60 in these two chlamydial species. Further studies are required to decipher these differences and understand the evolution of PGN-remodelling and division mechanisms in the *Chlamydiales* order.

Next, we raised antibodies against *S. negevensis* and *E. lausannensis* NlpC/P60 (Supplementary Fig. 14b,c) and investigated the subcellular localization of the proteins using confocal microscopy. In line with our observations in *C. trachomatis*, a peripheral cell envelope associated localization was observed in both *Chlamydia*-related genera throughout the developmental cycle (Fig. 9b,c). Contrasting with *C. trachomatis* where the YkfC protein was detected as punctae, the *S. negevensis* and *E. lausannensis* NlpC/P60 proteins were more evenly distributed over the envelope.

**Figure 9.**
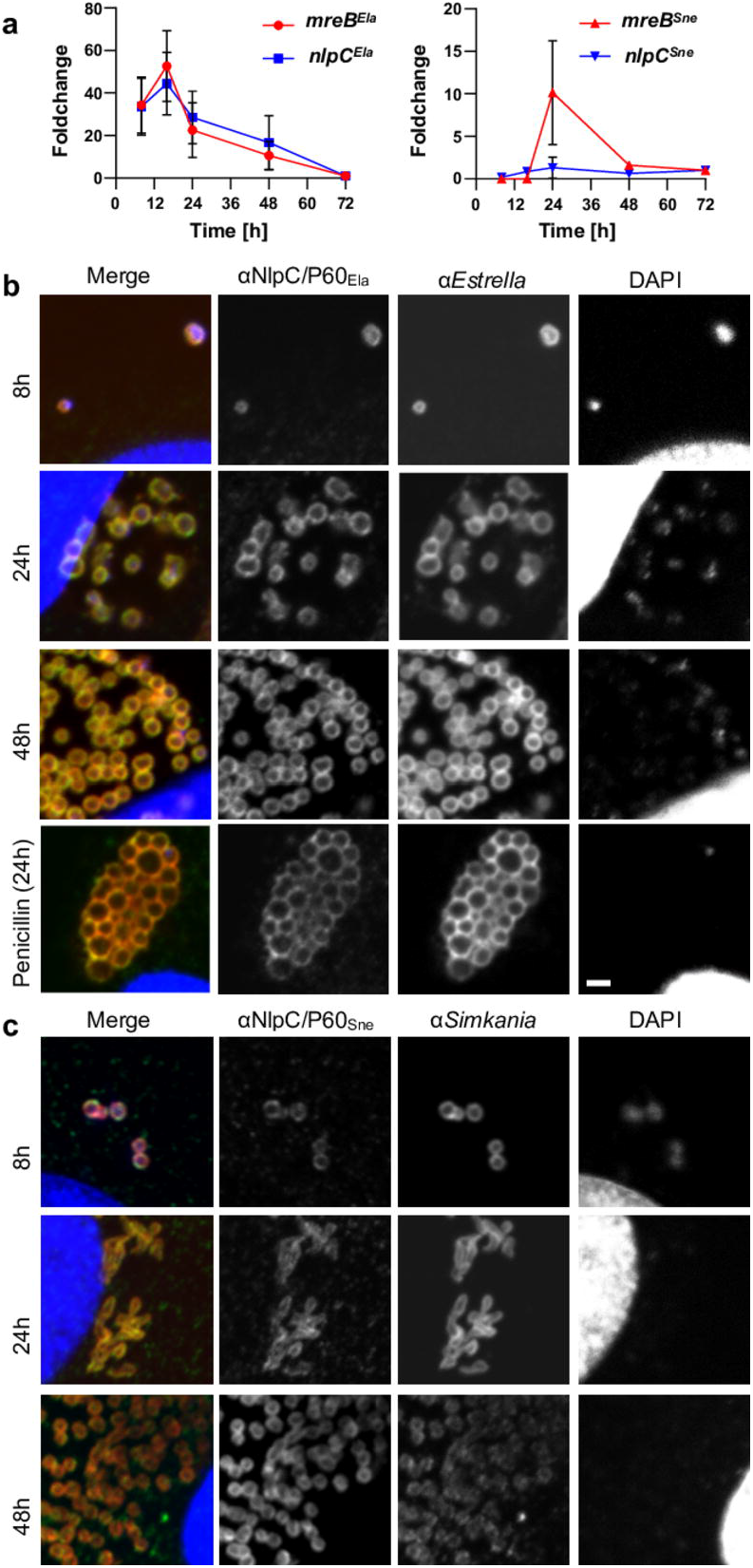
NlpC/P60 proteins of *Chlamydia*-related bacteria are expressed during the developmental cycle and show a peripheral localization. Vero cells infected with *S. negevensis* or *E. lausannensis* were harvested at the indicated times post-infection. (a) RNA levels of *nlpC* and *mreB* transcripts were quantified by qRT-PCR and normalized to the number of bacteria, which was estimated by qPCR (Supplementary Fig. 14a). RNA level is shown as foldchange compared to time 72 h. NlpC/P60_Ela_ (b) and NlpC/P60_Sne_ (c) show a peripheral localization throughout the developmental cycle of *E. lausannensis* and *S. negevensis*, respectively. Infected cells were harvested and labelled with anti-NlpC/P60 antibodies (green), species-specific antibodies (red) and DAPI (blue). Localization was investigated by confocal microscopy. Treatment with penicillin of *E. lausannensis* causes formation of enlarged aberrant bodies, but does not influence NlpC/P60 localization. Bars = 2 µm.

These divergent localization patterns may reflect potential differences in the organization and regulation of PGN machineries between *Chlamydia* and the evolutionary less reduced *Chlamydia*-related bacteria. Previous studies in chlamydiae indicated that persistence-inducing β-lactam antibiotics trigger delocalization of PGN biosynthesis enzymes [10]□. To test whether this effect also applies for PGN recycling enzymes, we treated *E. lausannensis* with penicillin but did not detect changes in the localization of the NlpC/P60 enzyme (Fig. 9b). *S. negevensis* were not investigated under penicillin treatment, as these chlamydiae are resistant to penicillin [45,46]. Taken together, we found NlpC/P60 proteins to be widespread among *Chlamydiales* and provide evidence that the homologs in the *Chlamydia*-related *S. negevensis* and *E. lausannensis*are were expressed throughout the developmental cycle and act as peripherally located PGN peptide recycling enzymes.

## Discussion

The paradoxical phenomenon of penicillin susceptibility in the absence of a PGN cell wall in *Chlamydiaceae* puzzled researchers for decades and has in the past been referred to as the chlamydial anomaly [47]□. The otherwise bactericidal antibiotic penicillin does not kill the intracellular pathogens but blocks cell division and induces reversible persistence [4,48,49]. In light of mechanistic insight into chlamydial PGN biosynthesis and the discovery of a PGN ring at the cell division plane, *Chlamydiaceae* emerged as model systems to understand how a minimal and modified PGN machinery supports bacterial cell division [4,5]. Synthesis of the transient PGN ring is likely tightly controlled and coordinated with the divisome to support robust chlamydial cell division. We are at the very early stages of understanding how chlamydial pathogens maintain a complete cycle of PGN ring synthesis, remodeling, and disassembly, including the process of PGN recycling.

Here, we present the first description of a chlamydial enzyme that catalyzes PGN recycling. The homolog of a *Bacillus* NlpC/P60 protein, YkfC_Ctr_, acts as a γ-D-Glu-mDAP peptidase that specifically recycles PGN-derived peptides in the human pathogen *C. trachomatis*. With the characterization of YkfC as a potentially important player in chlamydial PGN recycling we also contribute to our knowledge on the intimately linked processes of immune escape and energy recovery. Thereby, we identify the NlpC/P60 enzyme as a promising target for innovative anti-chlamydial drugs.

The evolutionary separated *Chlamydiaceae* do not harbor homologs of enzymes that recycle PGN-derived peptides in the model bacterium *E. coli*. Here, we found a member of the NlpC/P60 protein family to take over the essential step of decomposing PGN ring derived peptides in *C. trachomatis*. In the minimalist organism, the gene encoding *ykfC* (*ct127*) is not organized in a cluster of PGN peptide recycling genes as is known for its homolog from *Bacillu*s [50,51]. Instead, *ykfC* is located in close vicinity to genes involved in the uptake of glutamine, a process which was recently identified to be crucial for the initiation of chlamydial PGN synthesis and replication [52].

Chlamydiae feature an extraordinary biphasic developmental cycle. Elementary bodies (EBs) enter the host cell and differentiate into reticulate bodies (RBs) in a host derived vacuole. Exclusively in this intracellular stage, chlamydiae replicate, undergoing repeated cycles of cell division. Next, RBs undergo secondary differentiation into EBs, which are released from the host cell to re-start the developmental cycle. As expected for an enzyme which is involved in the cell division supporting PGN ring machinery, the encoding gene is transcribed during the mid-phase of the chlamydial developmental cycle when replication of RBs is occurring [40].

We propose a model in which YkfC has a central role in recycling of septal PGN ring material (Fig. 10). Tightly coordinated with constriction and daughter cell separation, the dual functioning enzymes SpoIID and AmiA degrade the PGN ring at mid-cell. Thereby, AmiA releases peptide side-chains from the PGN sugar backbone. This PGN processing step prevents recognition through innate immunity factor NOD2 which depends on PGN sugar unit MurNAc bound to the peptide side chain to percept chlamydial PGN. Nevertheless, the activity of AmiA leaves chlamydial cells at risk of detection by NOD1, another NOD receptor that senses Gram-negative PGN by recognizing the γ-D-Glu-mDAP dipeptide as its minimal ligand [4,42]. Our data help to answer the unsolved question how chlamydiae may impede NOD1 factor signaling. We hypothesize that peptides released by AmiA are transported from the periplasm to the cytoplasm through peptide permease OppABCDF [30] and then degraded by YkfC which specifically cuts the γ-D-Glu-mDAP bond. Thereby, YkfC breaks down the minimal peptidoglycan recognition motif of NOD1 and abolishes sensing of chlamydial infection - as suggested from our combined biochemical enzyme activity and *in vivo* NOD1 activation analyses. We speculate that OppABCDF and YkfC function together at the chlamydial cell membrane. This model is supported by our results from *C. trachomatis* and *Chlamydia*-related bacteria where NlpC/P60 proteins localized to the periphery of chlamydial cells. We propose that YkfC driven recycling of PGN-derived peptides does not exclusively contribute to pathogenicity. In parallel, it may close the cycle of PGN ring biosynthesis providing amino acids for lipid II precursor biosynthesis. Our biochemical and *E. coli* surrogate host derived data revealed that YfkC_Ctr_ releases mDAP which is an essential component of PGN that cannot be provided by the eukaryotic host cell. *De novo* synthesis of mDAP requires six energy-intensive biochemical reactions in *C. trachomatis* [4,53]. Thus, recycling of PGN-derived peptides is not only a result of chlamydial pathoadaptation but is also likely involved in energy recovery [5] and in maintaining PGN ring biosynthesis during cell division - in line with our observation of an overall enrichment of YkfC_Ctr_ near the division site as cell division progresses. Taken together, our data further support a model, in which the reduced-genome intracellular pathogens use a fine-tuned and energy-saving closed-loop PGN machinery for cell division and maintaining long-term residence in the infected host. Further research is needed to elucidate how other components besides the energy cost intensive mDAP might be recycled from chlamydial PGN-derived peptides.

**Figure 10.**
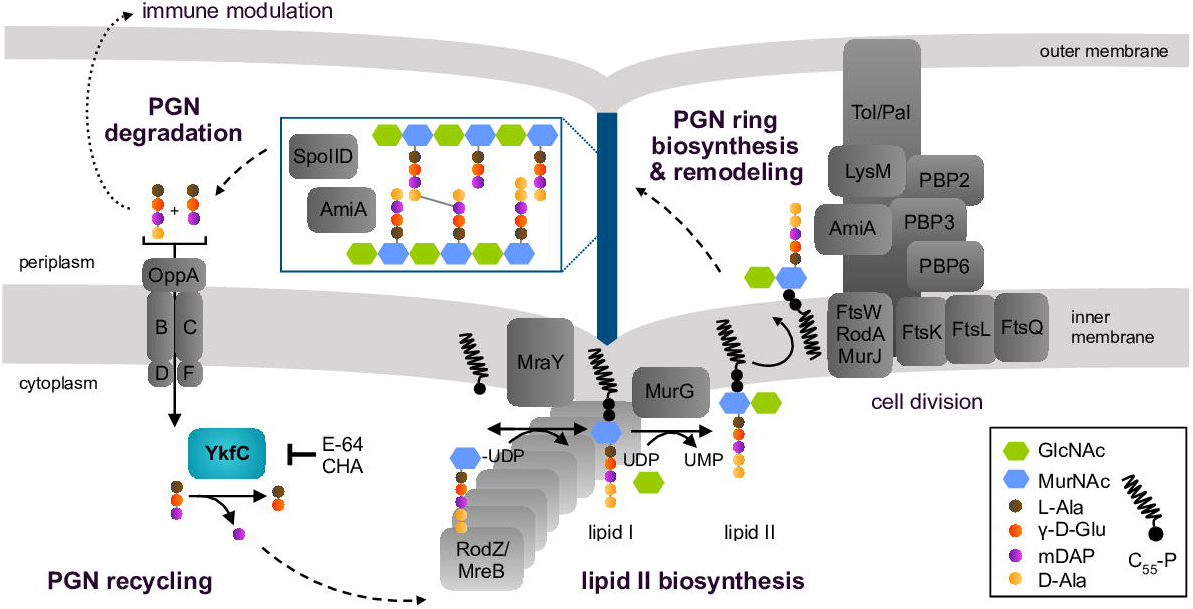
Proposed model for the PGN biosynthesis cycle in *C. trachomatis* that maintains metabolic robustness and cell division and modulates host immune response. PGN precursor lipid II is synthesized inside the chlamydial cell. The process is organized by actin-like protein MreB and RodZ in the absence of tubulin-like protein FtsZ. Upon translocation into the periplasm, the disaccharide-pentapeptide motif of lipid II is incorporated into the septal PGN ring by transpeptidases PBP2 and PBP3 and glycosyltransferases FtsW and RodA. Subsequently, different hydrolases including DD-carboxypeptidase PBP6 and the dual function enzymes SpoIID (lytic transglycosylase/muramidase) and AmiA (amidase/DD-carboxypeptidase) constantly remodel and break down the PGN ring to support cell division. PGN degradation products need to be recycled to (i) recover energy cost intensive intermediates, (ii) maintain a full cycle of PGN biosynthesis for cell division, and (iii) evade host response. AmiA releases peptide side chains from the sugar backbone and thereby prevents NOD2 perception. Peptide permease complex OppABCDF and NlpC/P60 peptidase YkfC_Ctr_ are proposed to form a functional unit to transport and directly recycle the released immunogenetic peptides. YkfC_Ctr_ cleaves between D-Glu and mDAP within PGN-derived peptides. Thereby, the peptidase likely abolishs recognition through NOD1 that depends on γ-D-Glu-mDAP as a minimal motif for sensing infection. In addition, YkfC_Ctr_-driven recycling of PGN-derived peptides makes mDAP available for another round of lipid II synthesis. The essential PGN component mDAP is energy cost intensive and cannot be provided by the host. We found YkfC_Ctr_ to be inhibited by model compounds chloroacetone and E-64 making the peptidase a promising antimicrobial target at the intersection of energy recovery, cell division and immune evasion. CHA: chloroacetone; LysM: homolog of LysM protein Cpn0902; GlcNAc: *N*-acetylglucosamine; MurNAc: *N*-acetylmuramic acid.

Functioning as a junction between pathogenicity and two central cellular processes, OppABCDF and YkfC might be essential for survival in obligate intracellular pathogens such as chlamydiae. In contrast to the peptide transporter, YkfC has a catalytic function making it a better candidate for antichlamydial drug development. In many bacterial pathogens, proteases and peptidases have vital roles in viability and pathogenicity and are considered untapped targets for antimicrobial drugs [54]□. YkfC combines features of classic antibiotic targets in cell viability and anti-virulence targets in pathogenicity. When studying model compounds, we found natural lead compound E-64 to target peptidase YkfC_Ctr_ *in vitro* and observed antichlamydial activity of its more membrane permeable variant E-64d. These data may make the peptidomimetic epoxide a promising scaffold for development of antichlamydial protease inhibitors.

## Material and Methods

### Bacterial strains and growth conditions

For propagation of *C. trachomatis* serovar L2, McCoy (kind gift of Dr. Harlan Caldwell, NIH) host cells were cultured at 37°C with 5% CO_2_ in Dulbecco’s Modified Eagle Medium (DMEM; Invitrogen, Waltham, MA) containing 10% (v/v) fetal bovine serum (FBS; Hyclone, Logan, UT) and 10 μg/mL gentamicin (Gibco, Waltham, MA). *C. trachomatis* serovar L2 (strain 434/Bu) lacking the endogenous plasmid (-pL2; kind gift of Dr. Ian Clarke, Univ. Southampton) was grown in McCoy cells for use in transformations. The infected McCoy cells were incubated in DMEM containing 10% FBS, 10 μg/mL gentamicin, 1 U/mL penicillin G, and 1 μg/mL cycloheximide. All cell cultures and *C. trachomatis* serovar L2 stocks were routinely tested for *Mycoplasma* contamination using the Mycoplasma PCR detection kit (Sigma, St. Louis, MO).

For propagation of *C. trachomatis* serovar D, HEp-2 host cells (ATCC CCL-23, Cell Lines Services) were cultured in DMEM supplemented with 10% (v/v) FBS, 1x MEM non essential amino acids solution, 1x MEM vitamin solution, 2.5 µg/mL amphotericin B and 50 µg/mL gentamicin (Gibco) at 37°C and 5% CO_2_. *C. trachomatis* D/UW-3/CX (ATCC VR-885) was grown in HEp-2 cells for use in MIC experiments. The infected cells were incubated in DMEM containing 10% (v/v) fetal calf serum (FCS), 1x MEM non essential amino acids solution, 1x MEM vitamin solution, 2.5 µg/mL amphotericin B, 50 µg/mL gentamicin and 1.2 µg/mL cycloheximide (Merck, Darmstadt, Germany). Cell culture was tested monthly for *Mycoplasma* contaminations using the Venor GeM OneStep Mycoplasma Detection Kit (Minerva Biolabs, Berlin, Germany). All MIC experiments were conducted in growth medium without cycloheximide, amphotericin and gentamicin.

All *E. coli* strains were maintained on Luria Bertani (LB) agar plates, unless stated otherwise. *E. coli* BL21(DE3) carrying YkfC_Ctr_ expression plasmids was maintained on agar plates containing 100 µg/mL ampicillin. For maintenance of the mutant strains, that were used in complementation experiments, the agar plates contained 200 µg/mL mDAP and 30 µg/mL chloramphenicol in case of the *E. coli* Δ*dapD*Δ*mpl* double mutant strain and 100 µg/mL kanamycin in case of *E. coli mepS* mutant strain. *E. coli* NEB10β (New England Biolabs, Ipswich, MA) cells were used for cloning and expansion of the *E*.*coli/C. trachomatis* shuttle vector. *E. coli* NEB10β was grown at 30°C in LB broth. All chemicals and antibiotics were obtained from Sigma unless otherwise noted.

### Cloning of *C. trachomatis*

From *C. trachomatis* serovar L2 (strain 434/Bu), we amplified the gene *ctl0382* (homolog of *ct127*) with a six histidine tag in the reverse primer using Phusion enzyme (NEB), and the PCR product was linked with pBOMBL::L2 linearized by EagI and KpnI using HiFi assembly system (NEB). The HiFi products were transformed into NEB10β competent cells (NEB) and plated on LB agar containing 100 μg/mL ampicillin.

### Immunofluorescence and confocal microscopy of *C. trachomatis*

McCoy cells were infected with the transformants, and the infected cells were either fixed with al-dehydes at 10.5 hpi or 100% MeOH for 10 min at 24 hpi after inducing expression of the construct at 4 hpi with 10nM aTc. The coverslips were stained with primary anti-body rabbit anti-6xH (Abcam, Cambridge, MA) and goat anti-major outer membrane protein (MOMP) of *C. trachomatis* (Meridian, Memphis, TN). Subsequently, the co-verslips were stained with secondary antibodies donkey anti-rabbit (594) and donkey anti-goat (488) (Invitrogen).

### Transformation into *C. trachomatis* without endogenous plasmid

A day before beginning the transformation, McCoy cells were plated in a six-well plate (1 × 10^6^ cells/well). *C. trachomatis* serovar L2 without plasmid (-pL2) (kind gift of Dr. Ian Clarke) was incubated with 2 μg plasmid in Tris-CaCl_2_ buffer (10 mM Tris-Cl pH 7.5, 50 mM CaCl_2_) for 30 min at room temperature. During this step, the McCoy cells were washed with 2 mL Hank’s Balanced Salt Solution (HBSS) media containing Ca^2+^ and Mg^2+^ (Gibco). After that, cells were infected with the transformants in 2 mL HBSS per well. The plate was centrifuged at 400 × g for 15 min at room temperature and incubated at 37°C for 15 min. The inoculum was aspirated, and DMEM containing 10% FBS and 10 μg/mL gentamicin was added per well. At 8 h post infection (hpi), media containing 1 or 2 U/mL penicillin G and 1 μg/mL cycloheximide was added to the wells after aspirating the initial post-infection media, and the plate was incubated at 37°C until 48 hpi. At 48 hpi, the transformants were harvested and used to infect new McCoy cells. These harvest and infection steps were repeated every 44-48 hpi until mature inclusions were observed (typically collected after two passages).

### Cloning of *E. coli*

For overproduction, the *ykfC* (*ct127*) gene from *C. trachomatis* D/UW-3/CX was amplified by PCR using the primers listed in Supplementary Table 1 and cloned into pET52b using the XmaI and EagI restriction sites to generate a protein with an N-terminal Strep-tag II followed by a human rhinovirus 3C protease cleavage site. For complementation assays, the *ykfC* genes from *C. trachomatis* D/UW-3/CX (*ct127*) and *C. pneumoniae* GiD (*cpn0245*) as well as the genes encoding the NlpC/P60 proteins of *E. lausannensis* CRIB 30 (ELAC_2173) and *S. negevensis* ATCC VR-1471/Z (SNE.A23160) were amplified by PCR using the primers listed in Supplementary Table 1 and cloned into pBAD24+.

### Site-directed mutagenesis

To generate active site mutants of the respective proteins, the residues C172 in YkfC_Ctr_, C173 in YkfC_Cpn_, C194 in NlpC/P60_Ela_ and C153 in in NlpC/P60_Sne_ were changed to alanine using the QuikChange Lightning Site-Directed Mutagenesis Kit (Agilent Technologies). The respective sense and antisense primers listed in Supplementary Table 1 were used according to the manufacturer’s instructions. Correct base changes were confirmed by sequencing.

### YkfC_Ctr_ overproduction and purification

For purification of YkfC_Ctr_, *E. coli* BL21(DE3) cells were transformed with pET52b-*ykfC*_*Ct*r_. Cells were grown in 2 liters TB autoinduction medium [56,57] at 37 ºC for 1 h. Temperature was shifted to 30 ºC and growth was continued for 20 h. Cells were harvested by centrifugation for 15 min at 7550 × *g* and 14 ºC and the resulting cell pellet was resuspended in 40 mL buffer A (25 mM MOPS pH 7.2, 1 M NaCl, 2 mM MgCl_2_) supplemented with 2 µg/mL polymyxin B (Carl Roth), 1 mM phenylmethyl sulfonyl fluoride (Merck) and 1 U/ml benzonase (Merck). Cells were broken by sonication, supplemented with 0.25 % CHAPS (Carl Roth) and further incubated at 4ºC for 1 h with gentle shaking. The suspension was centrifuged for 1 h at 310500 × *g* at 4ºC and the supernatant was poured onto a Strep-tactin column (1 ml bed volume) equilibrated with buffer A. After complete passage the column was washed 5 times with buffer B (25 mM MOPS pH 7.2, 500 mM NaCl, 2 mM MgCl_2_, 10% glycerol). The protein was eluted with buffer C (25 mM MOPS pH 7.2, 300 mM NaCl, 2 mM MgCl_2_, 10% glycerol and 50 mM biotin) and dialysed against 2 × 1 L dialysis buffer (MOPS pH 7.2, 150 mM NaCl, 2 mM MgCl_2_, 10% glycerol and 2 mM DTT) at 4 ºC overnight. Purity was determined by SDS-PAGE and aliquots stored at -80ºC. For inhibitor screens DTT was omitted in all buffers. The active site mutant YkfC_Ctr_C172A was overproduced and purified in the same manner as wild type YkfC_Ctr_.

### YkfC_Ctr_ activity assay

Endopeptidase assays were carried out in a final volume of 25 μl containing 50 mM MOPS pH 7.5, 150 mM NaCl, 2 mM MgCl_2_, 0.5 mM substrate and 2 µM enzyme. Each reaction was incubated at 37 ºC for 2 h and terminated by incubation for 5 min at 100 ºC. The reactions were centrifuged for 5 min at 17000 × *g* and 20 µl of the supernatant was analyzed by TLC on a HPTLC alumina silica gel 60 plate (Merck) using 2-butanol-pyridine-ammonia-water (39:34:10:26) as the mobile phase. Separation was visualized by ninhydrin staining. For lipid II as the substrate, reaction products were extracted with 20 µl of n-butanol/pyridine acetate (2:1 v/v, pH 4.2) and analyzed by TLC using chloroform-methanol-water-ammonia (88:48:10:1) as mobile phase and visualized by Hanessian’ s staining.

For PGN peptides derivedfrom *E. coli* or *B. subtilis* PGN as the substrate, PGN from each organism was treated with recombinant *B. subtilis* amidase CwlC as described [58]. CwlC derived peptides from *E. coli* or *B. subtilis* PGN were incubated with YkfC_Ctr_ and the reaction products were analyzed by HPLC-MS using an electrospray ionisation (ESI) time of flight mass spectrometer (MS) (microTOF II, Bruker Daltonics) connected with an UltiMate 3000 HPLC (Dionex). Soluble components in 5 µl samples were separated on a reversed-phase C18 column (Gemini 5 µm C18 110 Å 150 mm × 4.6 mm; Phenomenex) in an UltiMate 3000 HPLC (Dionex) by applying a 30-min linear gradient of 0–40% acetonitrile (buffer A: 100 % acetonitrile) at a flow rate of 0.2 ml/min. Before each run, the column was pre-equilibrated with 0.01% formic acid/0.05% ammonium formate, followed by the linear gradient to 100% acetonitrile. After each run, the column was rinsed for 12.45 min with 60% acetonitrile and subsequently re-equibrilated for 12.45 min with 0.01% formic acid/0.05% ammonium formate. The mass-to-charge rations of the separated samples were analysed by MS, operating in negative ion mode with a mass range of 120–1000.

Extracted ion chromatograms for the tripeptide (3P; theoretical mass [M−H]^−^ = 389.167), tetrapeptide (4P; [M−H]^−^ = 460.204), DAP ([M−H]^−^ = 189.087) and DAP-Ala ([M−H]^−^ = 260.124) were obtained within an error range of ± 0.02.

For PGN as the substrate, freshly prepared sacculi from *E. coli* W3110 were stained as described previously [20]□. The assay was carried out in a final volume of 100 µl containing 50 mM MOPS pH 7.5, 150 mM NaCl, 2 mM MgCl_2_, 1.25 µg sacculi and 4 µM YkfC_Ctr_. Each reaction was incubated for 16 h at 37°C. Insoluble material was removed by centrifugation for 10 min at 21000 × *g* and the supernatant containing soluble reaction products was measured on a spectrophotometer (Nano Photometer, Implen) at 595 nm.

### YkfC_Ctr_ inhibitor assays

All inhibitors were purchased from Merck Millipore (formerly Sigma Aldrich). The assay was carried out as described above with minor modifications. Before adding the substrate, YkfC_Ctr_ was pre-incubated with the inhibitor for 15 min at room temperature.

### *E. coli* Δ*mepS* complementation

The *E. coli* JW2163-3 Δ*mepS* mutant was received from the Keio *E. coli* mutant strain collection [59]. Competent cells of this *E. coli* mutant were transformed with the appropriate vectors (Supplementary Table 1). 5 ml of LB containing 100 μg/ml ampicillin were inoculated with material of a single, freshly transformed colony and incubated for 16 h at 37°C. For each construct material from three different colonies were tested separately in the assay. Subsequently, 500 µl of the culture was diluted 1/100 in nutrient broth (0.5% tryptone, 0.3% yeast extract) containing 100 μg/ml ampicillin. Cultures were incubated at 37°C and harvested at an OD_600_ of 1. Each cell pellet from 1 ml was suspended in 1 ml 0.9% NaCl solution and starting with this suspension a 1:10 dilution series in 0.9% NaCl was performed. The dilutions were plated on nutrient broth agar (NA) plates containing 0.04% L-arabinose using a replica plater (Merck). After drying the plates were incubated for 16 h at 30°C, 37°C and 42°C, respectively.

### Construction of the *E. coli* Δ*dapD*Δ*mpl* double mutant strain

The AT980 Δ*dapD* strain requiring mDAP for growth was earlier described [26]□. The MLD2502 strain derived from BW25113 that carries a deletion of the chromosomal *mpl* gene (∆*mpl*::Cm^R^) was described previously [60]. The ΔdapDΔ*mpl* double mutant strain used in this work was constructed by transduction of the ∆*mpl*::Cm^R^ mutation into the AT980 strain by phage P1 transduction [61]□.

### *E. coli* Δ*dapD* Δ*mpl* complementation

Competent cells of *E. coli* BW25113 Δ*dapD* Δ*mpl* were transformed with the appropriate vectors (Supplementary Table 1). 5 ml of LB containing 200 μg/ml mDAP and 100 μg/ml ampicillin were inoculated with material of a single, freshly transformed colony and incubated for 16 h at 37 °C. Subsequently, 500 µl of the culture was diluted 1/100 in TB medium [56] containing 200 μg/ml mDAP, 100 μg/ml ampicillin and 0.4 % L-arabinose. Cultures were incubated at 37 °C and harvested at an OD_600_ of 1. Each cell pellet from 1 ml was washed twice with 1 ml TB. The final suspension was diluted 1:3 in TB and for each well 100 µl were added to 100 µl TB containing 200 μg/ml ampicillin and 0.8% L-arabinose. Wells were further supplemented with 40 µM tripeptide. Samples were incubated for 6 h at 37 °C with intermittent shaking every 15 min. Growth was monitored at OD_600_ on a Tecan platereader.

### Immunofluorescence and confocal microscopy with *Chlamydia-related* bacteria

Vero cells (ATCC CCL-81) were cultivated in Dubelcco’s modified Eagle’s medium (DMEM) supplemented with 10% FBS at 37°C and 5% CO_2_. Cells were detached and 5 × 10^5^ cells/ml were incubated on a glass coverslip in 24 well plate overnight to allow them to adhere. *E. lausannensis* or *S. negevensis* cultivated in *Acanthameoba castellanii* were filtered with 5 μm filter to remove intact ameobae and diluted 1000 × in DMEM 10% FBS. Infection was synchronized by centrifugation for 15 min at 1790 × *g*. Infected cells were then incubated 15 min at 37°C, cells were washed and fresh medium was added.

After 24 h of incubation at 37 °C 5% CO_2_ infected cells were fixed with ice-cold meth-anol for 5 min, washed 3 times with PBS and incubated overnight in blocking solution (PBS, 1% BSA, 0.1% saponin). Immunofluorescence was performed as previously described [11]□. Coverslips were then incubated with antibodies: rabbit anti *E. lausannensis*, anti *S. negevensis* [62,63] and antibodies targeting NlpC of each bacterium (produced by immunization of mice with the purified proteins, Eurogentec, Seraing, Belgium). A subsequent incubation with secondary antibodies Alexa Fluor 488 goat anti-mouse and Alexa Fluor 594 donkey anti-rabbit (Thermo FisherScientific, (Waltham, MA) and DAPI was performed. Samples were then imaged by confocal microscopy (LSM 710 confocal microscopy, Zeiss, Oberkochen, Germany) and treated using ImageJ software (U. S. National Institutes of Health, Bethesda, MD).

### Transcription levels measurements

Cells were infected with *E. lausannensis* or with *S. negevensis* as described above and harvested by scrapping at the different time points post-infection. Samples were split and used for (i) quantification of the number of bacteria by qPCR, (ii) and gene expression measurement by qRT-PCR.

For qPCR, genomic DNA was isolated from 50 μl of cell suspension with the Wizard SV Genomic DNA purification system (Promega) following the manufacturer’s instructions, with a final elution in 200 μl of water. qPCR was performed on 5 μl of DNA and iTaq supermix with ROX (Bio-Rad). Primers SneF and SneR (400 nM each) and 200 nM of probe SneS were used to quantify genome copies of *S. negevensis* [63]. Primers EstF and EstR (400 nM each) and 200 nM of probe EstS were used to quantify *E. lausannensis* [62]. qPCR conditions were 5 min at 95°C followed by 40 cycles of 30 s at 95°C and 1 min at 60°C. Amplification and detection of PCR products were performed using a QuantStudio 3 (Applied Biosystems).

For qRT-PCR, RNA was stabilized from 500 μl of cell suspension using RNA Protect (Qiagen, Venlo, Netherlands) by incubation for 5 min at room temperature. Suspension was centrifuged at 5000 x *g* for 10 minutes, supernatant was removed, and pellet was kept at -20°C. RNA was extracted from the pellet using the RNeasy Plus Kit (Qiagen), following manufacturer’s instructions. DNA was eliminated by selective digestion with DNAse from Ambion DNA-free kit (Thermo Fisher Scientific, Waltham, MA). Reverse transcription was then performed to synthesize cDNA using a Goscript Reverse Transcription System (Promega). qPCR was used to quantify cDNA of *mre*B and *nlpC* for both *E. lausannensis* and *S. negevensis*. qPCR was performed by mixing 5 μl of 5-times diluted cDNA with 10 μl of iTaq Universal SYBR Green mix (Bio-Rad), 3.6 μl of water and 0.8 μl of forward and reverse primers listed in Supplementary Table 1. qPCR was performed using a QuantStudio 3 (Applied Biosystems, Waltham, MA) with the following conditions: 5 min of denaturation at 95°C followed by 45 cycles of 30 sec denaturation at 95°C and 1 min of annealing/elongation at 60°C.

### Minimal inhibitory concentration of cysteine protease inhibitors against *C. trachomatis*

The effect of cysteine protease inhibitors on active *C. trachomatis* infection was analysed by fluorescence microscopy. First, 3.5×10^4^ HEp-2 cells were seeded into black 96-well µ-plates (ibidi, Gräfelfing, Germany) and incubated for 3 days at 37°C and 5% CO_2_. Cells were washed once with medium and infected with *C. trachomatis* D/UW-3/CX. After 2 h the EB suspension was exchanged with fresh medium. The effect of E-64, chloroacetone (dissolved in water), and E-64d (dissolved in DMSO) on *C. trachomatis* was determined by addition of the inhibitors at 6, 10 and 24 hpi. Cells were fixed with 100% (v/v) icecold methanol at 30 hpi and washed once with 1× phosphate-buffered saline (PBS, 4 mM KH_2_PO_4_ pH 7.4, 16 mM Na_2_HPO_4_, 115 mM NaCl). Pathfinder chlamydia culture confirmation system (Bio-Rad Laboratories, Feldkirchen, Germany) was used for staining of *C. trachomatis* and host cell cytoplasm according to the manufacturers’ information. Genomic DNA was labeled with 10 µg/mL DAPI (Thermo Fischer Scientific, Waltham, MA) for 1 min and cells were washed twice with PBS for 10 min at 4 °C. Fluorescence microscopy was performed with an Axio observer Z.1 using Zen 2 software (Zeiss, Oberkochen, Germany).

### NOD1 reporter assay

Eukaryotic HEK-Blue hNOD1 cells (InvivoGen, Toulouse, France) were cultured as indicated in the manufacturers protocol. Briefly, cells were detached every 2-5 days by incubation with 1x PBS for 5 min at 37 °C. For passaging, the cell suspension was diluted with fresh DMEM supplemented with 10% (v/v) FBS, 2 mM L-glutamine, 100 µg/mL Normocin, 30 µg/mL blasticidin and 100 µg/mL Zeocin (InvivoGen, Toulouse, France) and incubated at 37°C and 5% CO_2_. The cell line was used to analyze release of NOD1 ligands by *C. trachomatis* during infection and treatment with cysteine protease inhibitors. 2×10^5^ HEK-Blue hNOD1 cells/mL were seeded into flat TC-96-well plates (Sarstedt, Nümbrecht, Germany) using supplemented DMEM lacking antibiotics and the plates were incubated for 20 h at 37°C and 5% CO_2_. Cells were washed once with medium and inoculated with *C. trachomatis* D/UW-3/CX at an infection rate of 20% for 2 h. Afterwards, the medium was exchanged with reconstituted HEK-Blue detection medium (InvivoGen, Toulouse, France). At 6 hpi, YkfC inhibitor chloroacetone (solved in H_2_O) was added in a serial dilution. Positive controls were conducted by the addition of 1.25 µg/mL triDAP 6 hpi (InvivoGen, Toulouse, France). Stimulation of NOD1 was measured at OD_600 nm_ 18 hpi and 24 hpi (corresponding to treatment durations of 12 h and 18 h, respectively) using a Tecan infinite M200 plate reader and Tecan SparkControl software (Tecan Group, Männedorf, Switzerland).

## Acknowledgments

Funding was provided by the Deutsche Forschungsgemeinschaft (DFG, German Research Foundation) Project-ID 398967434 —TRR261 and 390536577 and BONFOR, Medical Faculty, University of Bonn. J.R. and H.B. received a PhD fellowship from the Jürgen Manchot foundation. B.H. received support by the funding scheme FEMHABIL, Medical Faculty, University of Bonn. This project has also received funding from the European Union’s Horizon 2020 research and innovation program under the Marie Skłodowska-Curie Grant Agreement No. 721484 (International Training Network Train2Target) awarded to WV and was supported in part by the National Institutes of Health (NIH/NIGMS) through an award to SPO (1R35GM124798). We thank Katja Mölleken for providing us with genomic DNA from *C. pneumoniae*.

## Supporting information captions

## Supplementary Figures

**Supplementary Figure 1. Primary sequence alignment of YkfCBce, YkfCCtr, YkfCCpn and NlpC/P60Ela**.

**Supplementary Figure 2. Growth comparison of the E. coli** Δ**dapD**Δ**mpl double mutant containing either the empty vector or C. trachomatis YkfC constructs.**

**Supplementary Figure 3. Purification and in vitro activity of recombinant YkfCCtr**.

**Supplementary Figure 4. In vitro activity of YkfCCtr on B. subtilis tripeptide**.

**Supplementary Figure 5. In vitro activity of YkfCCtr on (non)PGN-derived peptides**.

**Supplementary Figure 6: TLC analysis of the inhibitory effect of chloroacetone on YkfCCtr activity at molar inhibitor:protein ratios below 8:1**.

**Supplementary Figure 7. Viability of HEp-2 cells upon treatment with cysteine protease inhibitors**.

**Supplementary Figure 8. Wild-type YkfCCtr expression levels are required for optimal chlamydial growth**.

**Supplementary Figure 9. Chlamydial inclusion formation is not affected by overexpression of YkfCCtr_6xH**.

**Supplementary Figure 10. Effect of cysteine protease inhibitor chloroacetone on the inclusion size of C. trachomatis exposed to concentrations below the MIC (4** μ**g/mL) and on uninfected HEK-Blue hNOD1 cells**.

**Supplementary Figure 11. Growth comparison of the E. coli** Δ**dapD**Δ**mpl double mutant containing either the empty vector or NlpC/P60 constructs. Supplementary**

**Figure 12. 3D in silico models of chlamydial NlpC/P60 enzymes. Supplementary**

**Figure 13. Primary sequence alignment of YkfCBce and NlpC/P60Sne**.

**Supplementary Figure 14. (a) Quantification of bacterial growth by 16S qPCR for E. lausannensis and S. negevensi. Antibodies raised against NlpC/P60 of E. lausannensis (b) and S. negevensis (c) specifically label bacteria**.

## Supplementary Material and Methods

**Cloning of C. trachomatis**.

**Immunofluorescence and confocal microscopy of C. trachomatis**.

**RT-qPCR analysis**.

**IFU assay. Cytotoxicity assay**.

## Supplementary Tables

**Supplementary Table 1. Strains, plasmids and oligonucleotides used in this study**.

