## Supplemental Information for "An NlpC/P60 protein catalyzes a key step in peptidoglycan recycling at the intersection of energy recovery, cell division and immune evasion in the intracellular pathogen *Chlamydia trachomatis*"

### Supplementary Information

#### Supplementary Figures

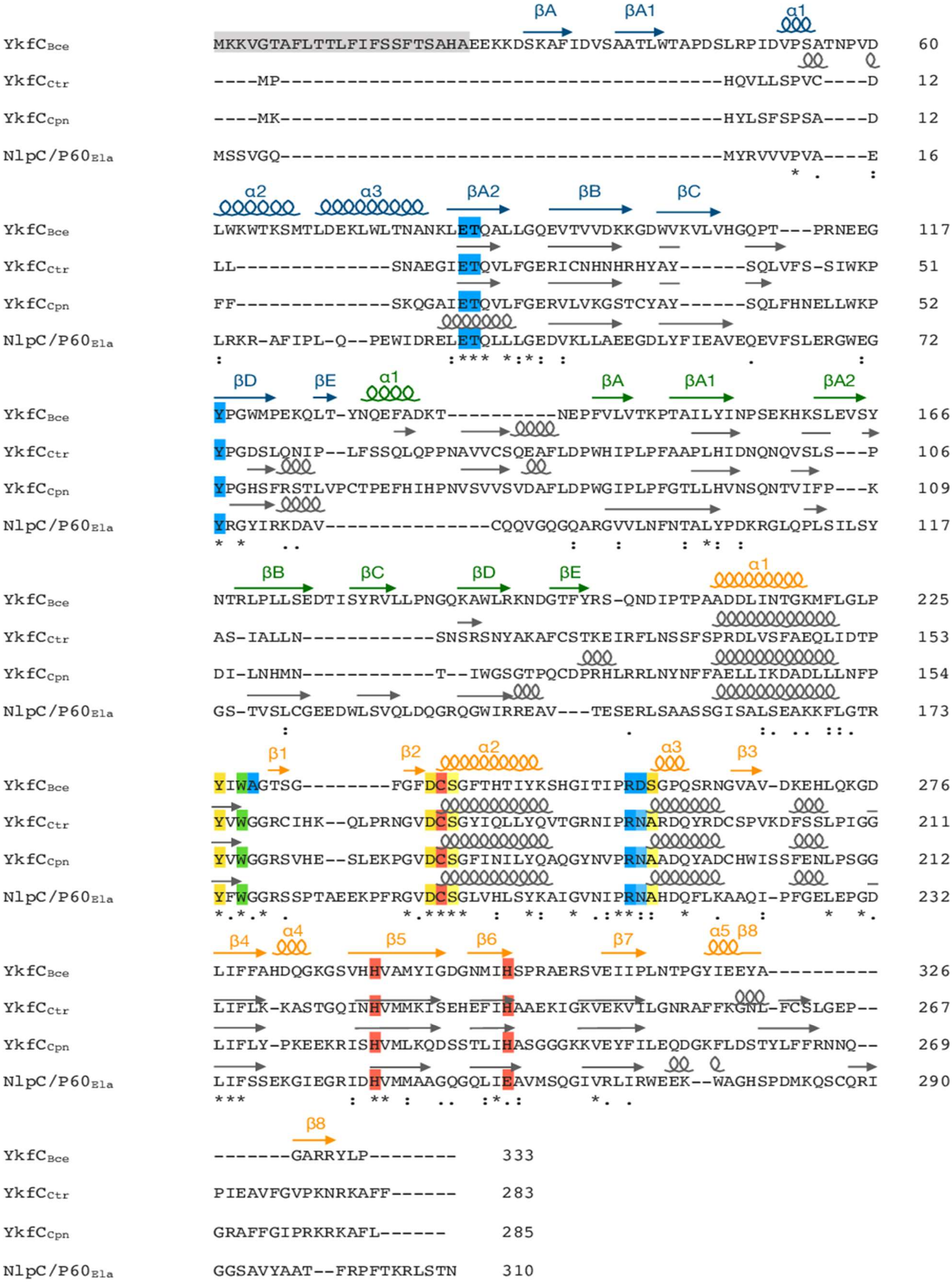

**Supplementary Figure 1. Primary sequence alignment of YkfC$_{Bce}$, YkfC$_{Ctr}$, YkfC$_{Cpn}$ and NlpC/P60$_{Ela}$.** The secondary structure of YkfC$_{Bce}$ characterized by Xu *et al.,* 2010 [1] is indicated above its sequence with the Sh3b1 domain in blue, the Sh3b2 domain in green and the NlpC/P60 domain in orange. The *in silico* predicted secondary structures of the chlamydial proteins are shown in grey above the respective sequences. The catalytic triad comprising Cys, His and a polar residue is shown in red, residues contributing to the S1 and S2 sites essential for substrate recognition in YkfC$_{Bce}$ are highlighted in yellow and blue, respectively. Residues contributing to both sites are further highlighted in green. The signal peptide sequence of YkfC$_{Bce}$ is displayed in grey.

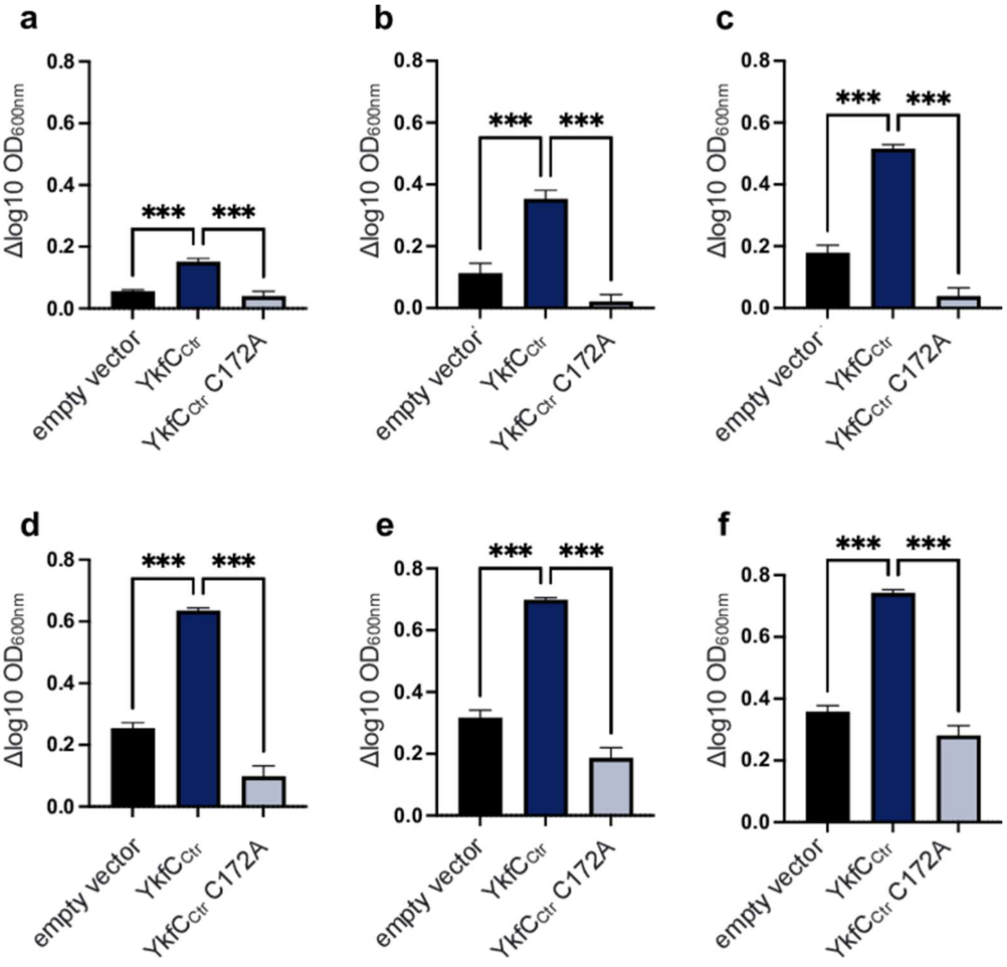


**Supplementary Figure 2. Growth comparison of *E. coli* Δ*dapD* Δ*mpl* double mutant containing either the empty vector or *C. trachomatis* YkfC constructs.** Growth at (a) 1 h, (b) 2 h, (c) 3 h, (d) 4 h, (e) 5 h and (f) 6 h was compared by one-way ANOVA analysis. P-values are shown as asterisks: n.s.: $P \geqslant 0.05$, *: P = 0.01 to 0.05, **: P = 0.001 to 0.01, ***: $P < 0.001$. Error bars indicate ± s.d.

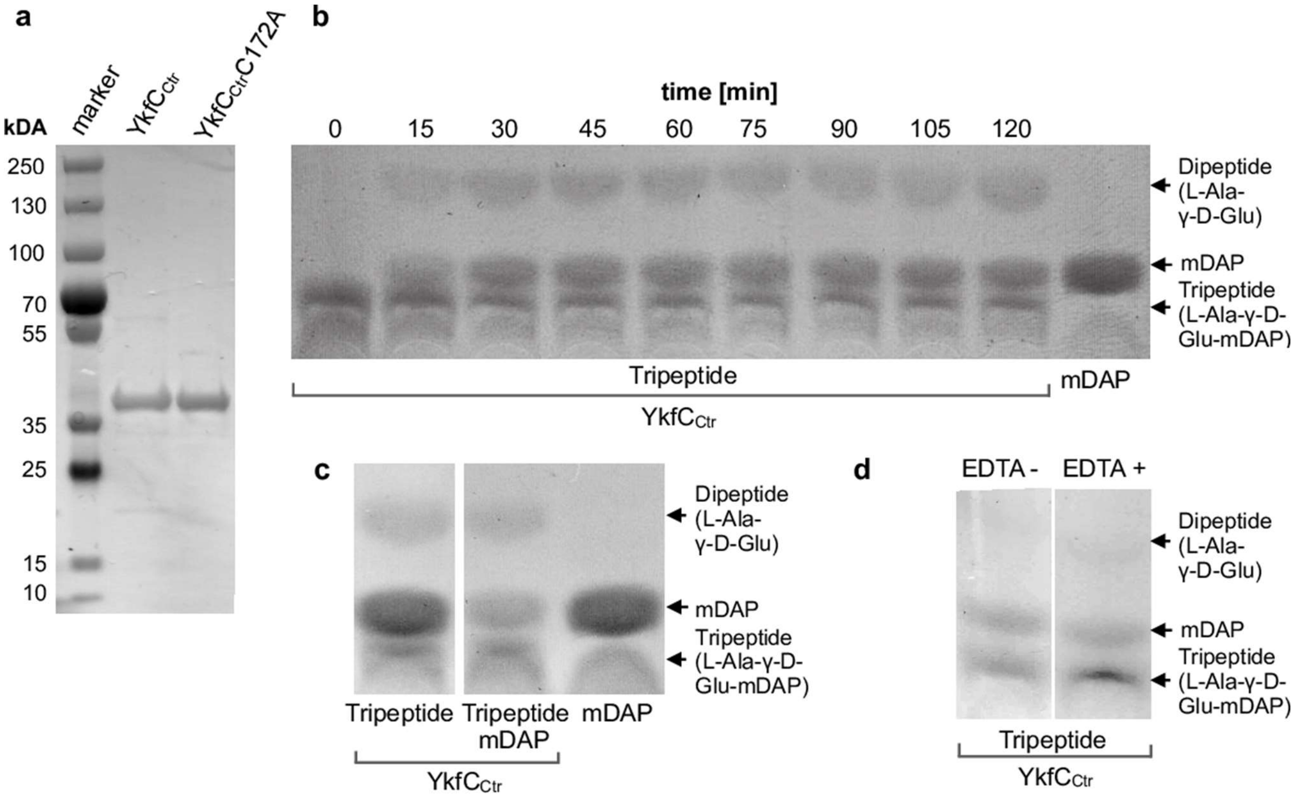


**Supplementary Figure 3. Purification and *in vitro* activity of recombinant YkfC$_{Ctr}$.** Recombinant YkfC$_{Ctr}$ as well as its active site mutant YkfC$_{Ctr}$C172A were analyzed by SDS-PAGE **(a)**. *In vitro* peptidase activity over time was analyzed by TLC using L-Ala-γ-D-Glu-mDAP tripeptide as substrate **(b)**. YkfC$_{Ctr}$ activity was time dependent reaching a plateau at t=30 min and was not decreased in presence of the reaction product mDAP (c) or the chelator EDTA **(d)**.

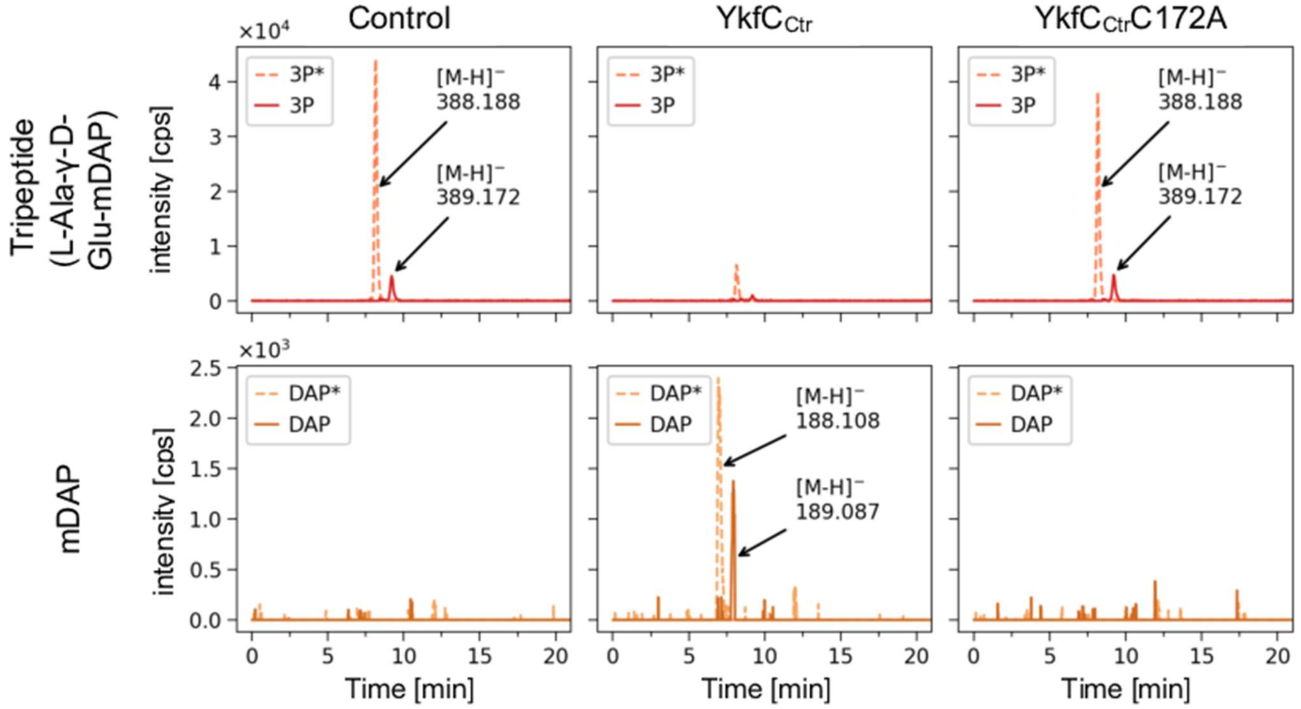


**Supplementary Figure 4. *In vitro* activity of YkfC$_{Ctr}$ on *B. subtilis* tripeptide.** LC-MS analysis of YkfC$_{Ctr}$ activity on CwlC derived amidated tripeptide from *B. subtilis* PGN. YkfC$_{Ctr}$, but not its active-site mutant YkfC$_{Ctr}$C172A, was active on this substrate regardless of the amidation of the free carboxyl group in mDAP as shown by the release of amidated mDAP. Data presented as extracted ion chromatograms for theoretical masses of amidated tripeptide (3P*, $[M-H]^{-}$ = 388.183), tripeptide (3P, $[M-H]^{-}$ = 389.167), amidated DAP (DAP*, $[M-H]^{-}$ = 188.103) and DAP ($[M-H]^{-}$ = 189.087) within an error of ± 0.02.

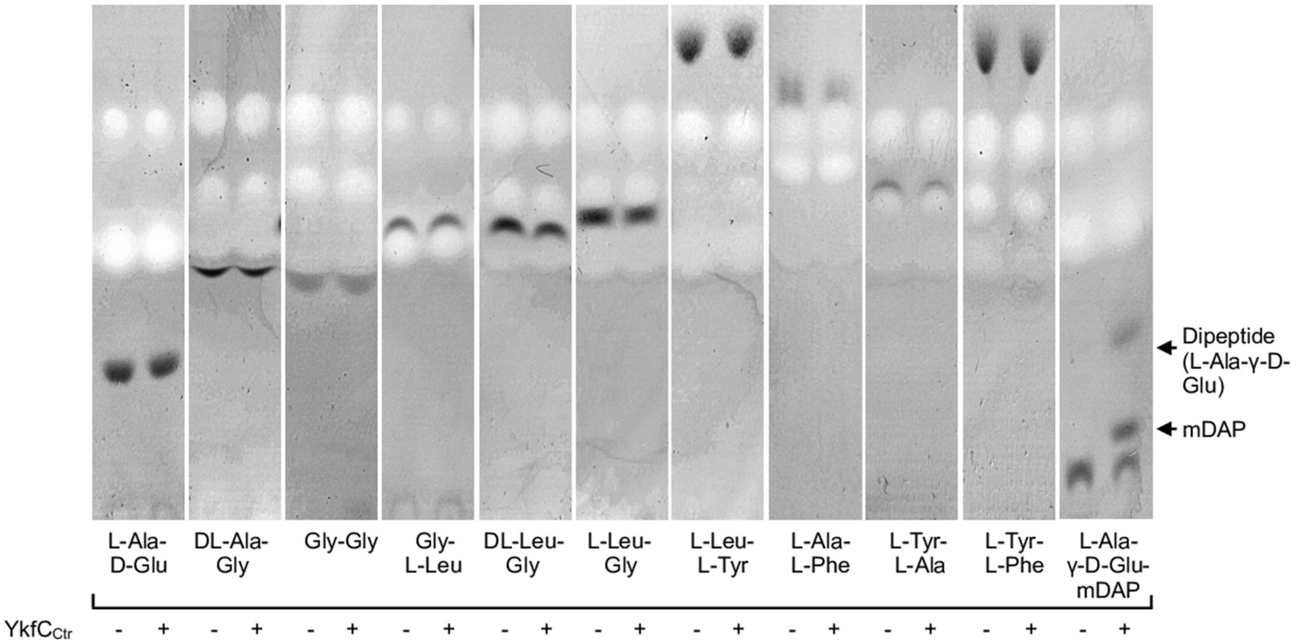


**Supplementary Figure 5. *In vitro* activity of YkfC$_{Ctr}$ on (non)-PGN derived peptides.** *In vitro* peptidase activity towards 10 (non)-PGN-derived peptides was analyzed by TLC. None of the peptides was used as asubstrate by YkfCCtr. L-Ala-γ-D-Glu-mDAP was used as positive control.

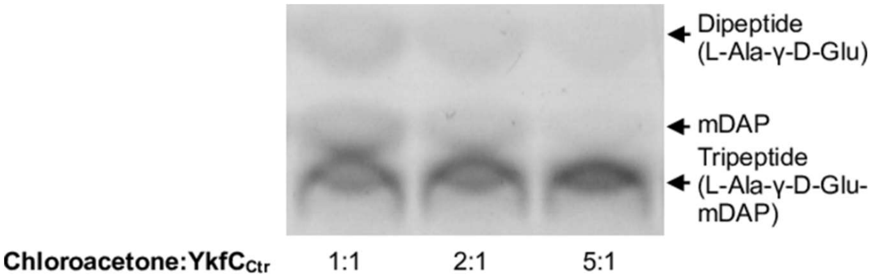


**Supplementary Figure 6. TLC analysis of the inhibitory effect of chloroacetone on YkfC$_{Ctr}$ activity at molar inhibitor:protein ratios below 8:1** (corresponding to a molar ratio inhibitor:active site cysteine of 1:1, provided that the additional seven cysteines in YkfC$_{Ctr}$ are accessible for the alkylating chemical). YkfC$_{Ctr}$ retained activity at all tested inhibitor:protein ratios below 8:1 as shown by the presence of reaction product mDAP.

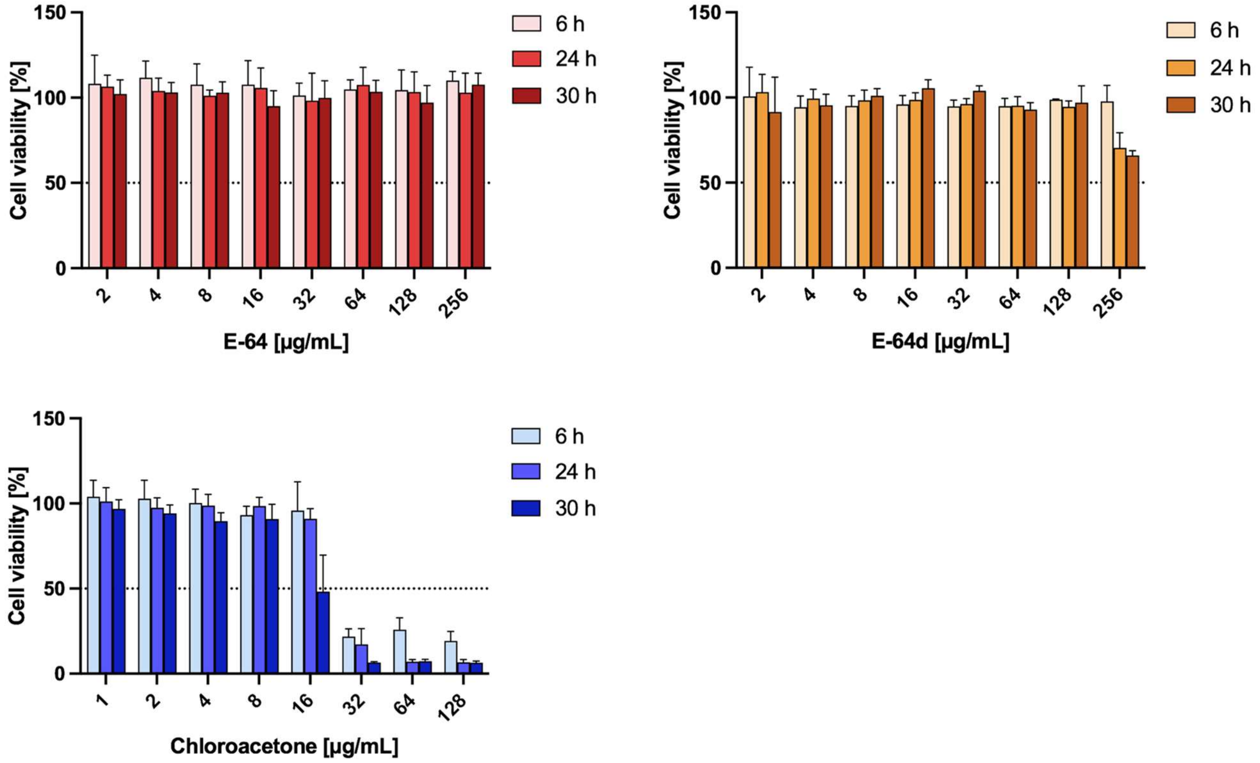


**Supplementary Figure 7. Viability of HEp-2 cells upon treatment with cysteine protease inhibitors.** Inhibitors were tested on confluent HEp-2 monolayer for 6 h, 24 h or 30 h. Cytotoxicity was visualized using alamarBlue cell viability reagent. Error bars indicate ± s.d. (n = 3).

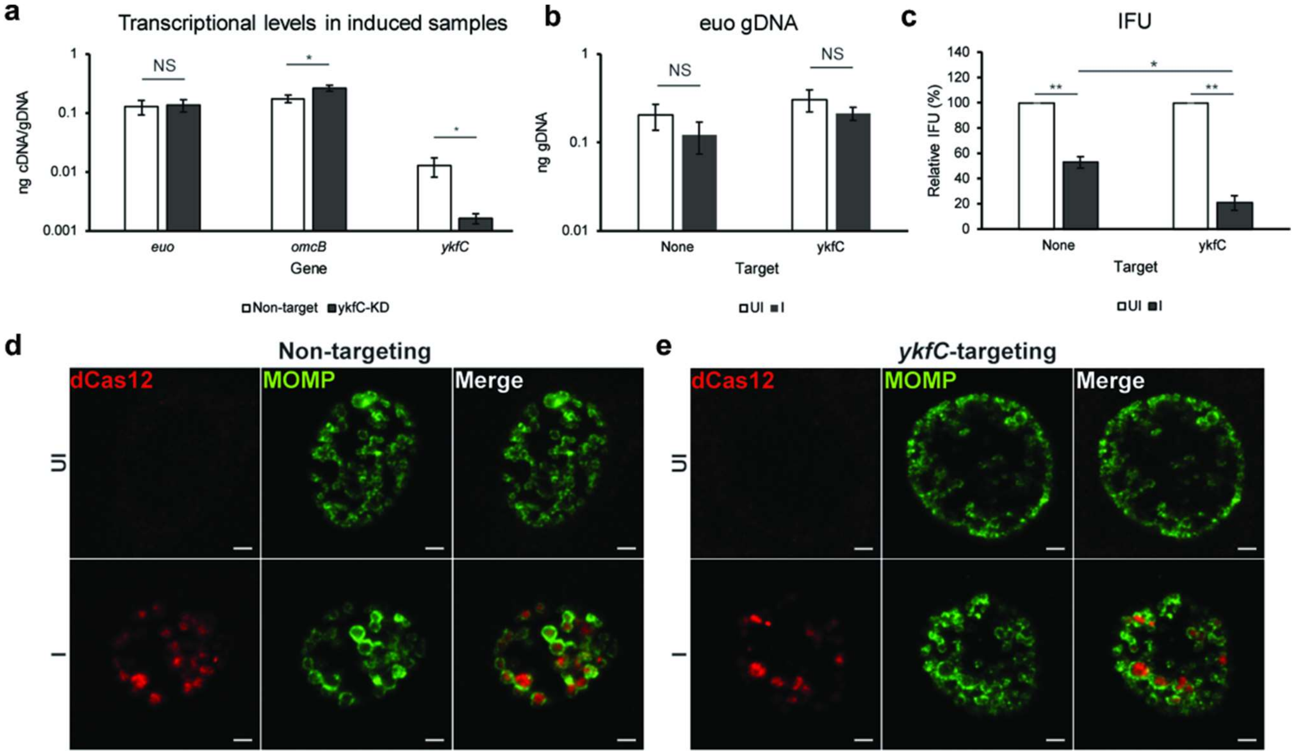


**Supplementary Figure 8. Wild-type YkfC$_{Ctr}$ expression levels are required for optimal chlamydial growth.** Constructs encoding dCas12-crRNA CRISPRi system targeting *ykfC$_{Ctr}$* or a non-targeting control were transformed into *C. trachomatis* (-pL2). McCoy cells were infected with these transformants, and dCas12 expression was induced or not with 10 nM aTc at 4 hpi. At 24 hpi, RNA and genomic DNA were isolated and used for (RT)-qPCR to measure transcript and genomic DNA levels. **(a)** Transcript levels of *euo*, *omcB*, and *ykfC* in dCas12-induced conditions at 24 hpi in the non-targeting or *ykfC$_{Ctr}$*-targeting knockdown strains. **(b)** The levels of genomic DNA of non-targeting and *ykfC$_{Ctr}$*-targeting knockdown strains at 24 hpi in uninduced (UI) or induced (I) conditions. **(c)** Inclusion forming units of non-targeting and *ykfC$_{Ctr}$*-targeting knockdown strains at 24 hpi in uninduced and induced conditions. **(a-c)** Error bars represent ± standard deviation (n = 3). *: $p<0.05$; **: $p<0.001$ **(d, e)** IFA controls of non-targeting and *ykfC$_{Ctr}$*-targeting knockdown strains at 24 hpi. MOMP (green) and dCas12 (red) were stained. The images were acquired on a Zeiss AxioImager.Z2 equipped with an Apotome2 using a 100X objective. Scale bar: 2 µm.

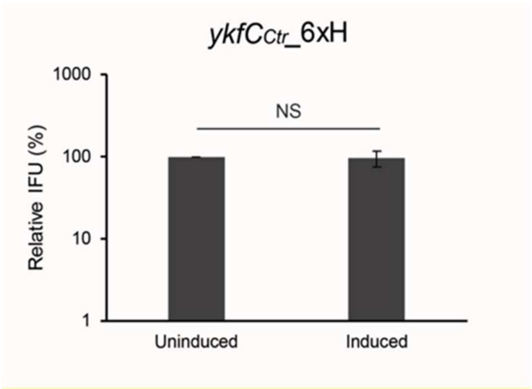


**Supplementary Figure 9. Chlamydial inclusion formation is not affected by overexpression of YkfC$_{Ctr}$_6xH**. The effect of YkfC$_{Ctr}$_6xH overexpression on inclusion formation was tested by inclusion forming unit (IFU) assay. McCoy cells were infected with uninduced and induced IFU samples. At 24 hpi, the infected cells were fixed, stained with goat anti-MOMP, and the inclusion number was counted. NS: not significant.

**a**

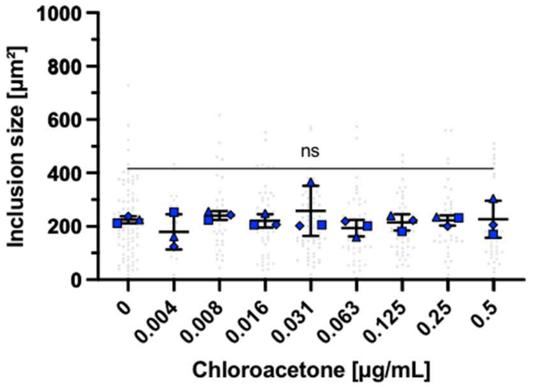


**b**

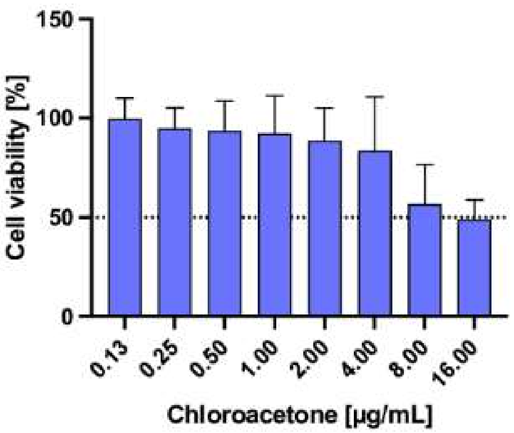


**c**

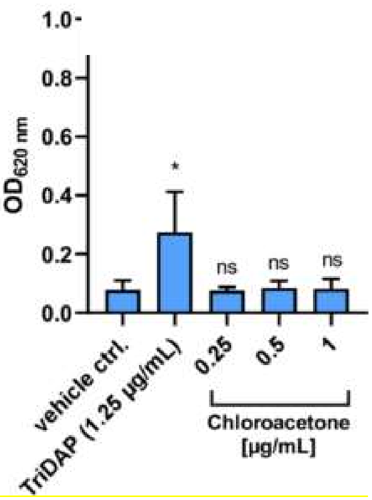


**Supplementary Figure 10. Effect of cysteine protease inhibitor chloroacetone on the inclusion size of *C. trachomatis* exposed to concentrations below the MIC (4 µg/mL) and on uninfected HEK-Blue hNOD1 cells. (a)** Effect on inclusion size. Chloroacetone did not impact on inclusion size in the tested sub-MIC range of concentrations (0.004-0,5 µg/mL) including concentrations that exerted an increase in NOD1 activation (0.016-0,031 µg/mL). **(b)** Effect on cell viability**.** Chloroacetone was tested in a concentration range of 0.13-16 µg/mL for 24 h treatment duration on uninfected HEK-Blue hNOD1 cells. Cytotoxicity was analyzed using alamarBlue cell viability reagent. Error bars indicate ± s.d. (n=4). **(c)** Effect on NOD1 stimulation. Chloroacetone was tested on uninfected HEK-Blue hNOD1 cells in a concentration range of 0.25-1 µg/mL for 18h. Stimulation of NOD1 was measured at $OD_{620\ nm}$. TriDAP was used as positive control. Error bars indicate ± s.d. (n=3).

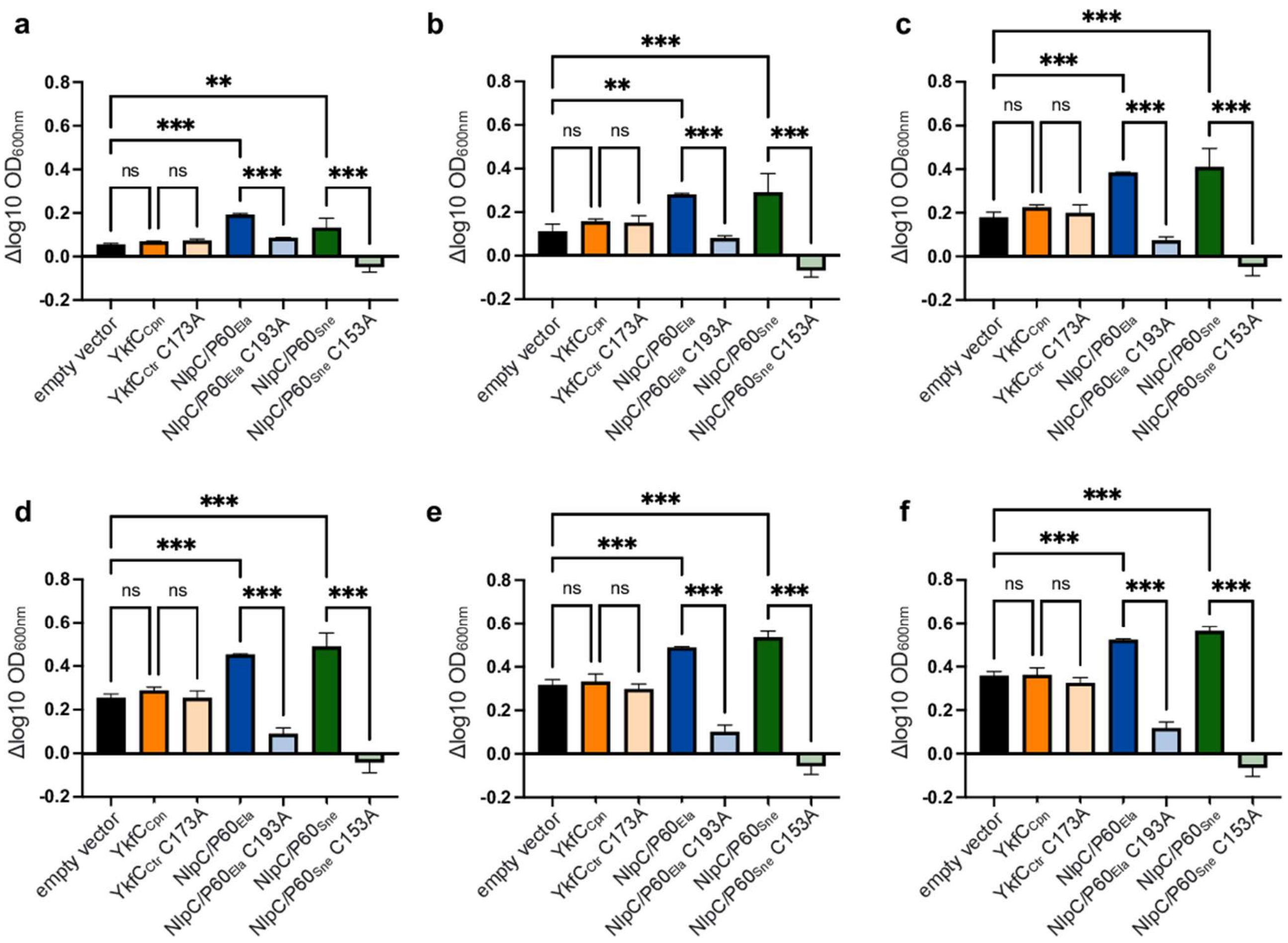


**Supplementary Figure 11. Growth comparison of *E. coli* Δ*dapD*Δ*mpl* double mutant containing either the empty vector or chlamydial NlpC/P60 constructs.** Growth at (a) 1 h, (b) 2 h, (c) 3 h, (d) 4 h, (e) 5 h and (f) 6 h was compared by one-way ANOVA analysis. P-values are shown as asterisks: n.s.: $P \geqslant 0.05$, *: P = 0.01 to 0.05, **: P = 0.001 to 0.01, ***: $P < 0.001$. Error bars indicate ± s.d.

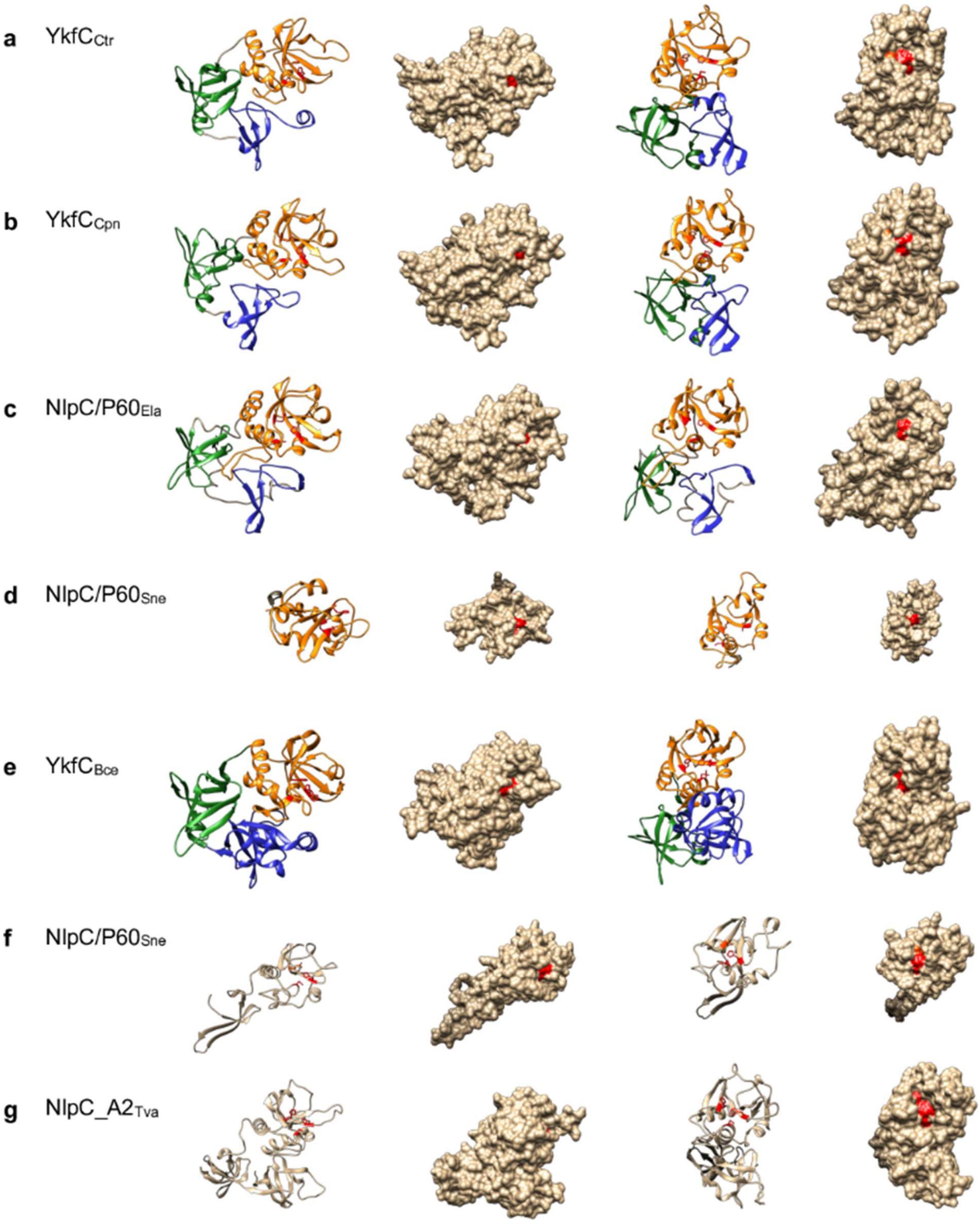


**Supplementary Figure 12. 3D *in silico* models of chlamydial NlpC/P60 enzymes.** The 3D structures of YkfC$_{Ctr}$ (**a**), YkfC$_{Cpn}$ (**b**), NlpC/P60$_{Ela}$ (**c**) and NlpC/P60$_{Sne}$ (**d**) were modeled based on Phyre2 alignment against the structure of YkfC$_{Bce}$ (**e**) characterized by Xu *et al.*, 2010 [1]. The Sh3b1 domain is highlighted in blue, the Sh3b2 domain in green and the NlpC/P60 domain in orange. The catalytic triad comprising Cys, His and a polar residue is shown in red. For NlpC/P60$_{Sne}$ another 3 D structure was predicted based on Phyre2 alignment against the structure of *Trichomonas vaginalis* NlpC/P60 protein NlpCA2 (NlpC_A2$_{Tva}$) characterized by Pinheiro *et al.*, 2019 [2] because this protein showed highest sequence and secondary structure matches in the database with *Simkania* NlpC/P60 (**f,g**).

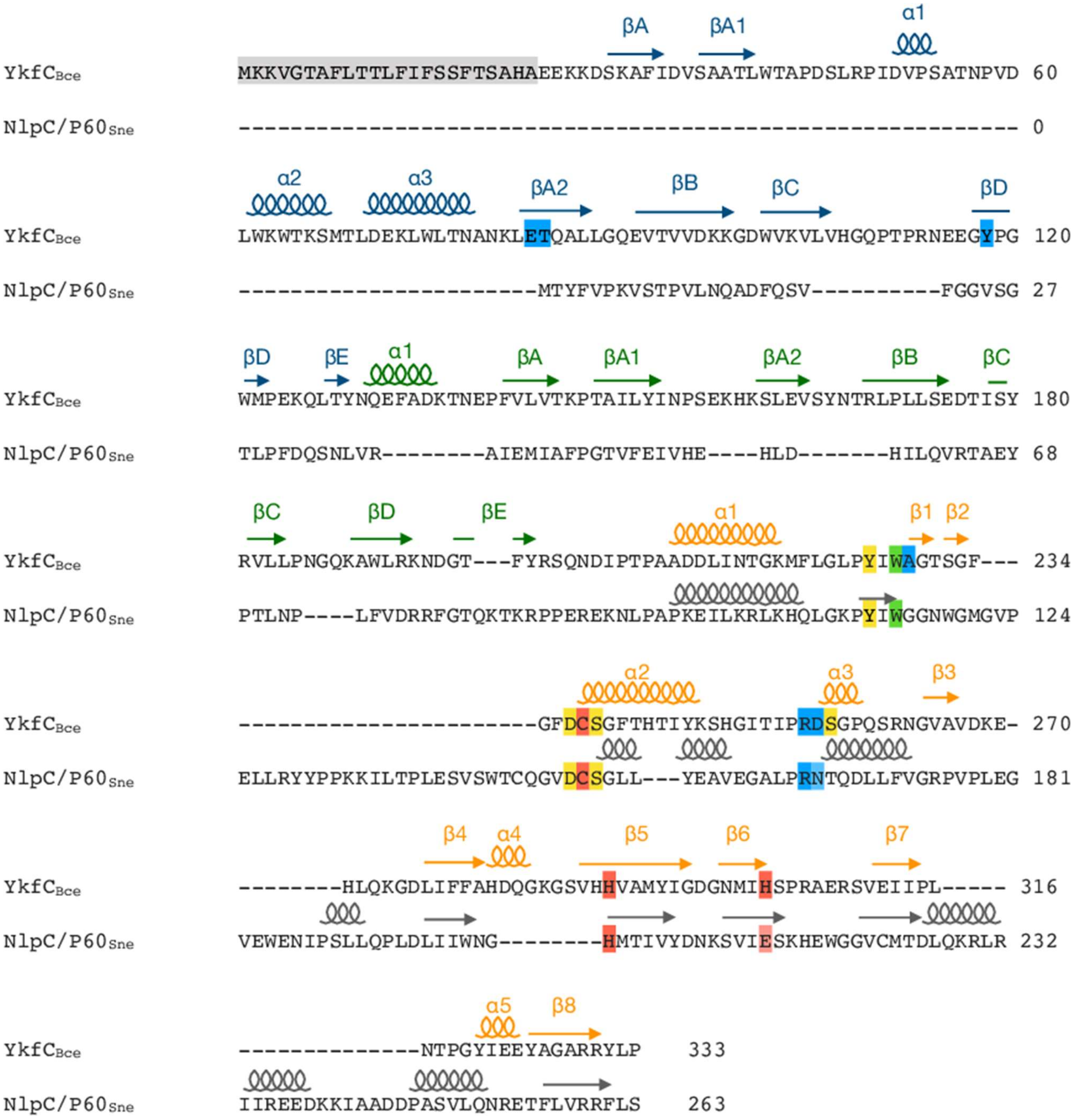


**Supplementary Figure 13. Primary sequence alignment of YkfC$_{Bce}$ and NlpC/P60$_{Sne}$.** The secondary structure of YkfC$_{Bce}$ characterized by Xu *et al*., 2010 [1] is indicated above its sequence with the Sh3b1 domain in blue, the Sh3b2 domain in green and the NlpC/P60 domain in orange. The in silico predicted secondary structure of NlpC/P60$_{Sne}$ is shown in grey above the respective sequences. The catalytic triad comprising Cys, His and a polar residue is shown in red, residues contributing to the S1 and S2 sites essential for substrate recognition in YkfC$_{Bce}$ are highlighted in yellow and blue, respectively. Residues contributing to both sites are further highlighted in green. The signal peptide sequence of YkfC$_{Bce}$ is displayed in grey.

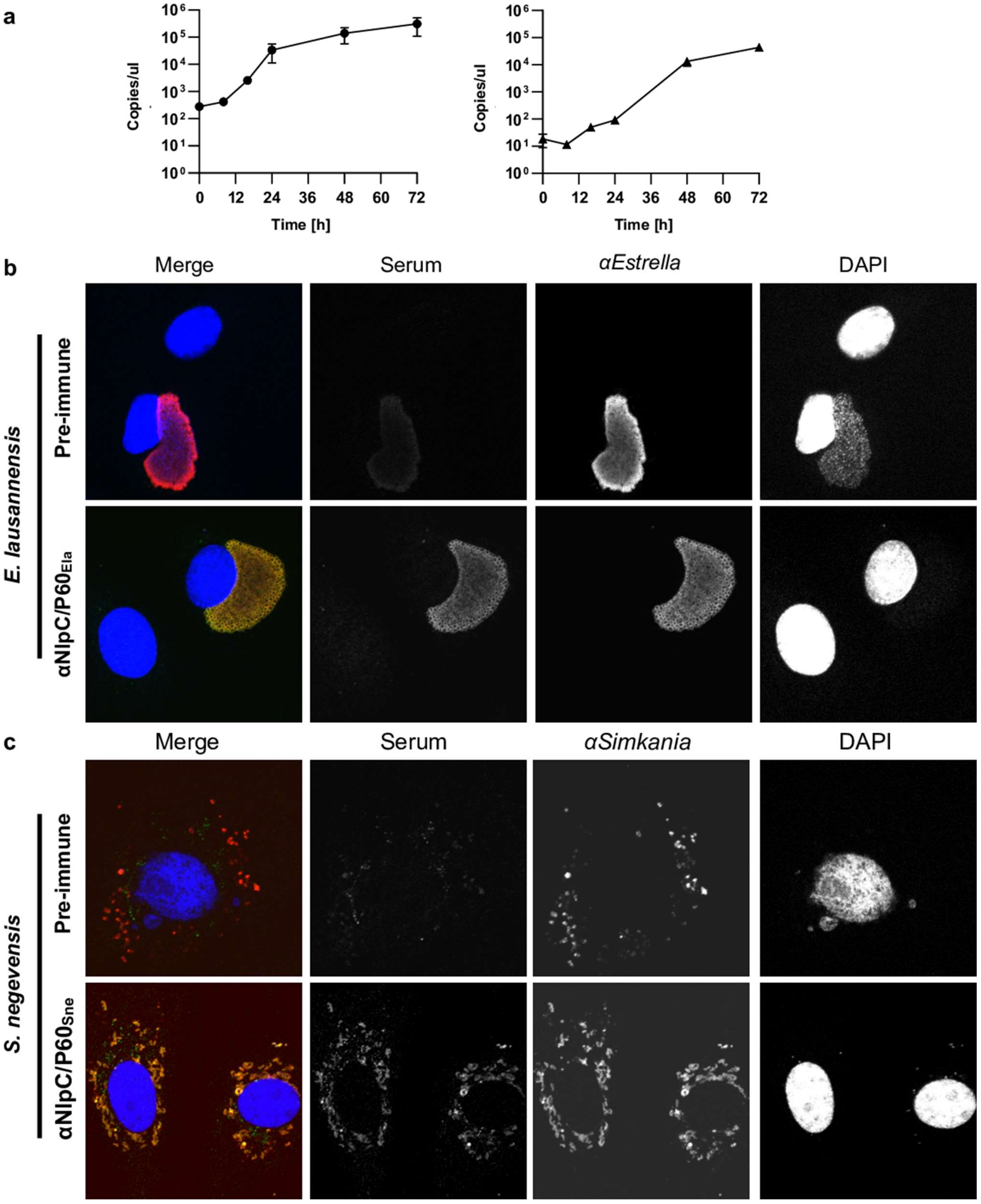


**Supplementary Figure 14. (a) Quantification of bacterial growth by 16S qPCR for *E. lausannensis* and *S. negevensis*.** DNA was extracted and copy number of bacterial genomes was quantified by qPCR targeting the 16S gene. This quantification was used to normalize the RNA quantification according to the number of bacteria in the sample. **Antibodies raised against NlpC/P60 of *E. lausannensis* (b) and *S. negevensis* (c) specifically label bacteria.** Cells infected with *S. negevensis* or *E. lausannensis* were labelled with mice serum taken before (pre-immune) or after immunization with the indicated protein (green), with species-specific antibodies (red) and DAPI (blue). Bar = 2 µm.

#### Supplementary Material and Methods

**Cloning of *C. trachomatis*.** For *ctl0328* (homolog of *ct127*)-knockdown in *C. trachomatis* serovar L2 (strain 434/Bu), a gBlock (IDTDNA, Coralville, IA) encoding a crRNA cassette was designed to target the 5‘ intergenic region and inserted into pBOMBL12CRia linearized by BamHI using HiFi (NEB) as described [3]. As a control, a scrambled non-targeting crRNA gBlock cassette was also inserted into the BamHI-digested pBOMBL12CRia empty vector. The HiFi products were transformed into NEB10β competent cells (NEB) and plated on LB agar containing 100 µg/mL ampicillin.

**Immunofluorescence and confocal microscopy of *C. trachomatis***. For IFA controls of qPCR, McCoy cells were infected with the transformants, and the infected cells were fixed with 100% MeOH for 10 min. The coverslips were stained with primary antibody goat anti-major outer membrane protein (MOMP) of *C. trachomatis* (Meridian, Memphis, TN) and mouse anti-dCas12 (Millipore-Sigma). Subsequently, the coverslips were stained with secondary antibodies donkey anti-goat (488) or donkey anti-mouse (594) (Invitrogen).

**RT-qPCR analysis.** McCoy cells were infected with the *C. trachomatis* L2 transformants carrying the aTc-inducible constructs encoding dCas12 with or without constitutively expressed crRNA targeting $ykfC_{Ctr}$ or the non-targeting control. At 4hpi, the constructs were induced or not with 10 nM aTc. RNA and genomic DNA were collected using Trizol (Invitrogen) and DNeasy Tissue (Qiagen) kit as described previously [4]. The isolated RNA was treated with DNase to remove DNA contamination, and cDNA was synthesized with the DNased RNA using Superscript III reverse transcriptase (Invitrogen). Equal volumes of cDNA were used in qPCR reactions [4]. Similarly, equal mass of DNA was used from each sample in qPCR reactions using the primers listed in Supplementary Table 1 to quantify chlamydial genomes, which were used to normalize the amount of cDNA.

**IFU assay.** The transformants were infected into McCoy cell monolayers in 6-well plate (MOI=0.1). At 4 hpi, the contructs were induced with 0 or 10 nM aTc, and the transformants were harvest at 24 hpi and frozen at -80°C. The IFU samples were thawed and diluted to $10^{-4}$. The diluted cells were infected into McCoy cells in 24-well plates. At 24 hpi, the infected cells were fixed and stained with goat anti-MOMP (594), and the number of inclusions was counted.

**Cytotoxicity assay.** Cytotoxicity of cysteine protease inhibitors on HEp-2 and HEK-Blue hNOD1 cells, was assessed via the resazurin-based alamarBlue cell viability reagent (Thermo Fisher Scientific, Waltham, MA). $5x10^4$ HEp-2 cells/mL or $2x10^5$ HEK-Blue hNOD1 cells/mL were seeded into flat TC-96-well plates (Sarstedt, Nümbrecht, Germany) and incubated 2 days for HEp-2 and 20 h for HEK-Blue hNOD1 cells at 37 °C and 5% $CO_2$. Next, cells were washed with medium followed by the addition of inhibitors in a serial dilution within the indicated concentration range. After 6, 24 or 30 h incubation with the compound, cells were washed twice with Hanks' balanced salt solution (HBSS) and incubated with alamarBlue cell viability reagent diluted 1:10 in HBSS for 1 h. Convertion of resazurin into resorufin by viable cells was determined by transferring the supernatant into black 96-well plates (Greiner Bio-One, Frickenhausen, Germany) and measuring the fluorescence at 550 nm excitation and 595 nm emission wavelength using a Tecan infinite M200 plate reader and Tecan SparkControl software (Tecan Group, Männedorf, Switzerland).

#### Supplementary Tables

**Supplementary Table 1. Strains, plasmids and oligonucleotides[a] used in this study.**

| Strain, plasmid or oligonucleotide | Description | Reference or source |
|---|---|---|
| *E. coli* BL21(DE3) | Expression strain | New England Biolabs |
| *E. coli* JW2163-3 | *ΔmepS* mutant, used for complementation assays | Keio collection; Baba et al., 2006 [5] |
| *E. coli* Δ*dapD* Δ*mpl* | *ΔmplΔdapD* double mutant, used for complementation assays | this study |
| *C. trachomatis* D/UW-3/CX (ATCC VR-885) | *C. trachomatis* strain used for MIC and hNOD1 activation experiments | ATCC |
| HEp-2 cells (ATCC CCL-23) | Used for propagation of *C. trachomatis* in MIC experiments | ATCC |
| HEK-Blue hNOD1 cells | Used for hNOD1 assay with *C. trachomatis* | InvivoGen, Toulouse, France |
| Vero cells (ATCC CCL-81) | Used for propagation of *Chlamydia*-like organisms | ATCC |
| pET52b-ykfC$_{Ctr}$ | *ykfC* from *C. trachomatis*, N-terminal Strep-tag, used for cytoplasmic overproduction | this study |
| pET52b-ykfC$_{Ctr}$C172A | *ykfC* from *C. trachomatis*, N-terminal Strep-tag, YkfC$_{Ctr}$ C172A mutant (YkfC$_{Ctr}$C172A), used for cytoplasmic overproduction | this study |
| pBAD24+-ykfC$_{Ctr}$ | *ykfC* from *C. trachomatis*, used for complementation assays | this study |
| pBAD24+ykfC$_{Ctr}$C172A | *ykfC* from *C. trachomatis*, YkfC$_{Ctr}$ C172A mutant (YkfC$_{Ctr}$C172A), used for complementation assays | this study |
| pBAD24+-ykfC$_{Cpn}$ | *ykfC* from *C. pneumoniae*, used for complementation assays | this study |
| pBAD24+ykfC$_{Cpn}$C173A | *ykfC* from *C. pneumoniae*, YkfC$_{Cpn}$ C173A mutant (YkfC$_{Cpn}$C173A), used for complementation assays | this study |
| pBAD24+-nlpC/P60$_{Ela}$ | *nlpC/P60* from *E. lausannensis*, used for complementation assays | this study |
| pBAD24+-nlpC/P60$_{Ela}$C193A | *nlpC/P60* from *E. lausannensis*, NlpC/P60$_{Ela}$ C193A mutant (YkfC$_{Ela}$C172A), used for complementation assays | this study |
| pBAD24+-nlpC/P60$_{Sne}$ | *nlpC/P60* from *S. negevensis*, used for complementation assays | this study |
| pBAD24+-nlpC/P60$_{Sne}$C153A | *nlpC/P60* from *S. negevensis*, NlpC/P60$_{Sne}$ C153A mutant (YkfC$_{Sne}$C153A), used for complementation assays | this study |
| pBAD24+-mepS$_{Eco}$ | *mepS* from *E. coli*, used for complementation assays | this study |
| ykfC$_{Ctr}$-pET52_fow | gcgcgcCCCGGGccgcaccaagtcttatt | this study |
| ykfC$_{Ctr}$-pET52_rev | gcgcgcCGGCCGtcaaaagaaggcttttctatttttag | this study |
| ykfC$_{Ctr}$-pBAD24+_fow | GAGGAATTCACCATGCCGCACCAAGTCTTATTGTCTCCT | this study |
| ykfC$_{Ctr}$-pBAD24+_rev | TCATCCGCCAAAACATCAAAAGAAGGCTTTTCTATTTTT | this study |

| | | |
|---|---|---|
| ykfC$_{Cpn}$-pBAD24+_fow | GAGGAATTCACCATGAAACACTACCTATCATTTTCTCCT | this study |
| ykfC$_{Cpn}$-pBAD24+_rev | TCATCCGCCAAAACATTACAGAAAGGCTTTTCTTTTTCT | this study |
| nlpC/P60$_{Ela}$-pBAD24+_fow | GAGGAATTCACCATGAGTTCTGTCGGGCAGATGT | this study |
| nlpC/P60$_{Ela}$-pBAD24+_rev | TCATCCGCCAAAACATCAGTTGGTGGACAGTCTCTT | this study |
| nlpC/P60$_{Sne}$-pBAD24+_fow | GAGGAATTCACCATGACCTATTTTGTCCCAAAAGTTTCAA | this study |
| nlpC/P60$_{Sne}$-pBAD24+_rev | TCATCCGCCAAAACATTACGAGAGGAATCGTCGGACT | this study |
| mepS$_{Eco}$-pBAD24+_fow | GAG GAA TTC ACC ATG GTC AAA TCT CAA CCG ATT TTG AG | this study |
| mepS$_{Eco}$-pBAD24+_rev | TCA TCC GCC AAA ACA TTA GCT GCG GCT GAG AAC CC | this study |
| ykfC$_{Ctr}$-C172A_fow | acagcttcctcgtaatggtgtagatgcttcggggtatattc | this study |
| ykfC$_{Ctr}$-C172A_rev | gaatataccccgaagcatctacaccattacgaggaagctgt | this study |
| ykfC$_{Cpn}$-C173A_fow | agtctggaaaagccgggtgttgatgcttcgggatttatca | this study |
| ykfC$_{Cpn}$-C173A_rev | tgataaatcccgaagcatcaacacccggcttttccagact | this study |
| nlpC/P60$_{Ela}$-C193A_fow | gcacaagaccggaagcatccactcccctgaaagg | this study |
| nlpC/P60$_{Ela}$-C193A_rev | cctttcaggggagtggatgcttccggtcttgtgc | this study |
| nlpC/P60$_{Sne}$-C153A_fow | ggacctgtcaaggagtggatgcctcgggacttctt | this study |
| nlpC/P60$_{Sne}$-C153A_rev | aagaagtcccgaggcatccactccttgacaggtcc | this study |
| nlpC(ct127)/qPCR F | AACCATCGACACTATGCCTATTC | this study |
| nlpC(ct127)/qPCR R | TGAGAGCAGACAACAGCATTAG | this study |
| nlpC(ct127)/(pBOMBL)/F | aaagatcttcacacaggacatctgcATGCCGCACCAAGTCTTATTATC | this study |
| nlpC(ct127)_6xH/(pBOMBL)/R | acatatttgaatggtcgaccggtacttaatggtgatggtgatggtgAAAGAAGGCTTTTCTATTTTTAGGAAC | this study |
| nlpC(ct127) crRNA gBlock | tgtgaaagtgggtcttaagacgtcggtactgcatgtgacgcacgtagatcatgcaTTCACCGGTGGAGACGGTTTTCTTATAATGACACCTAATTTCTACTCTTGTAGATGTCCTCTTTGTGATGTCTCTACAAATAAAACGAAAGGCTCAGTCGAAAGACTGGGCCTTTCGTTTTATcaacagcggtctactgaatctgagctagtgcgtgatataattaaaattatattca | this study |
| non-targeting crRNA gBlock | tgtgaaagtgggtcttaagacgtcggtactgcatgtgacgcacgtagatcatgcaTTCACCGGTGGAGACGGTTTTCTTATAATGACACCTAATTTCTACTCTTGTAGATACCGAGTTGCCCGTTAAAGTACAAATAAAACGAAAGGCTCAGTCGAAAGACTGGGCCTTTCGTTTTATcaacagcggtctactgaatctgagctagtgcgtgatataattaaaattatattca | this study |
| $^{sne}$NlpC_F | GTTCCAGAACTTTTACGTTA | this study |
| $^{sne}$NlpC_R | AAAGAGCAAATCTTGTGTAT | this study |
| $^{ela}$NlpC_F | AGTTACAAAGCAATAGGAGT | this study |
| $^{ela}$NlpC_R | CATCATCACATGATCAATC | this study |
| $^{sne}$MreB_F | GTAAAACACCTCGTAAGATT | this study |

| | | |
|---|---|---|
| [sne]MreB_R | GTACAGCAATTAAAATTTTG | this study |
| [ela]MreB_F | TTAAGGCTCTGATCAAAC | this study |
| [ela]MreB_R | CGATCAAAATAACCTCTT | this study |

[a] in 5‘-3‘ direction.
